# Analytical kinetic model of native tandem promoters in *E. coli*

**DOI:** 10.1101/2021.11.19.468961

**Authors:** Vatsala Chauhan, Mohamed N.M. Bahrudeen, Cristina S.D. Palma, Ines S.C. Baptista, Bilena L.B. Almeida, Suchintak Dash, Vinodh Kandavalli, Andre S. Ribeiro

**Affiliations:** Laboratory of Biosystem Dynamics, Faculty of Medicine and Health Technology, Tampere University, Finland; Department of Cell and Molecular Biology, Uppsala University, Uppsala, Sweden

## Abstract

Closely spaced promoters in tandem formation are abundant in bacteria. We investigated the evolutionary conservation, biological functions, and the RNA and single-cell protein expression of genes regulated by tandem promoters in *E. coli*. We also studied the sequence (distance between transcription start sites ‘*d_TSS_’*, pause sequences, and distances from oriC) and potential influence of the input transcription factors of these promoters. From this, we propose an analytical model of gene expression based on measured expression dynamics, where RNAP-promoter occupancy times and *d_TSS_* are the key regulators of transcription interference due to TSS occlusion by RNAP at one of the promoters (when *d_TSS_* ≤ 35 bp) and RNAP occupancy of the downstream promoter (when *d_TSS_* > 35 bp). Occlusion and downstream promoter occupancy are modeled as linear functions of occupancy time, while the influence of *d_TSS_ i*s implemented by a continuous step function, fit to *in vivo* data on mean single-cell protein numbers of 30 natural genes controlled by tandem promoters. The best-fitting step is at 35 bp, matching the length of DNA occupied by RNAP in the open complex formation. This model accurately predicts the squared coefficient of variation and skewness of the natural single-cell protein numbers as a function of *d_TSS_*. Additional predictions suggest that promoters in tandem formation can cover a wide range of transcription dynamics within realistic intervals of parameter values. By accurately capturing the dynamics of these promoters, this model can be helpful to predict the dynamics of new promoters and contribute to the expansion of the repertoire of expression dynamics available to synthetic genetic constructs.

**Author Summary:** Tandem promoters are common in nature, but investigations on their dynamics have so far largely relied on synthetic constructs. Thus, their regulation and potentially unique dynamics remain unexplored. We first performed a comprehensive exploration of the conservation of genes regulated by these promoters in *E. coli* and the properties of their input transcription factors. We then measured protein and RNA levels expressed by 30 *Escherichia coli* tandem promoters, to establish an analytical model of the expression dynamics of genes controlled by such promoters. We show that start site occlusion and downstream RNAP occupancy can be realistically captured by a model with RNAP binding affinity, the time length of open complex formation, and the nucleotide distance between transcription start sites. This study contributes to a better understanding of the unique dynamics tandem promoters can bring to the dynamics of gene networks and will assist in their use in synthetic genetic circuits.

## Introduction

Closely spaced promoters exist in all branches of life in convergent, divergent, and tandem formations [1–7]. Models of tandem promoters [8–10] have largely been based on measurements of synthetic constructs [11–13] and predict that such promoter arrangements result in unique transcription dynamics due to the interference between RNAPs transcribing the promoters [9, 10, 14–19].

When an RNAP is committed to form the open complex (OC), a process lasting up to hundreds of seconds [20–22], it occupies approximately 35 base pairs (bp), from the transcription start site (TSS, position 0) until position -35 [23–25]. If the TSS of a neighbouring promoter is closer than 35 bp away, it will not be possible for both promoters to be occupied simultaneously, since an RNAP occupying one of them will ‘occlude’ the other, preventing it from being reached [9]. However, if the promoters are more than 35 bp apart, this occlusion does not occur. Instead, interference will occur when RNAPs elongating from the upstream promoter collide with an RNAP occupying the downstream promoter [14] (in either closed or open complex formation), forcing one of the RNAPs to fall-off (both scenarios are likely possible, and we expect it to differ with, e.g., the binding affinity of the RNAP to the downstream promoter). Meanwhile, models based on empirical parameter values suggest that collisions between two elongating RNAPs are rare (because events such as pausing or simultaneous initiations from both promoters are rare). Also, even if and when such collisions occur, they are unlikely to result in fall-offs since the RNAPs are moving at similar speeds and in the same direction [9][10][26].

Models suggest that both forms of interference decrease the mean RNA production rate while increasing its noise based on the distance between promoters (*d_TSS_*), their strengths [10], and the time spent between commitment of the RNAP to OC and escape from the promoter region [27]. These hypotheses have yet to be empirically validated in natural tandem promoters.

We studied how *d_TSS_* and the time spent by RNAPs on the TSSs affect gene expression dynamics due to interference between the transcription processes of tandem promoters (Fig 1). We consider only the natural tandem promoters that neither overlap with nor have in between another gene (positionings I and II, which differ in if the promoter regions overlap or not) (see the other arrangements in Fig. S1 in the S2 Appendix).

**Fig 1.**
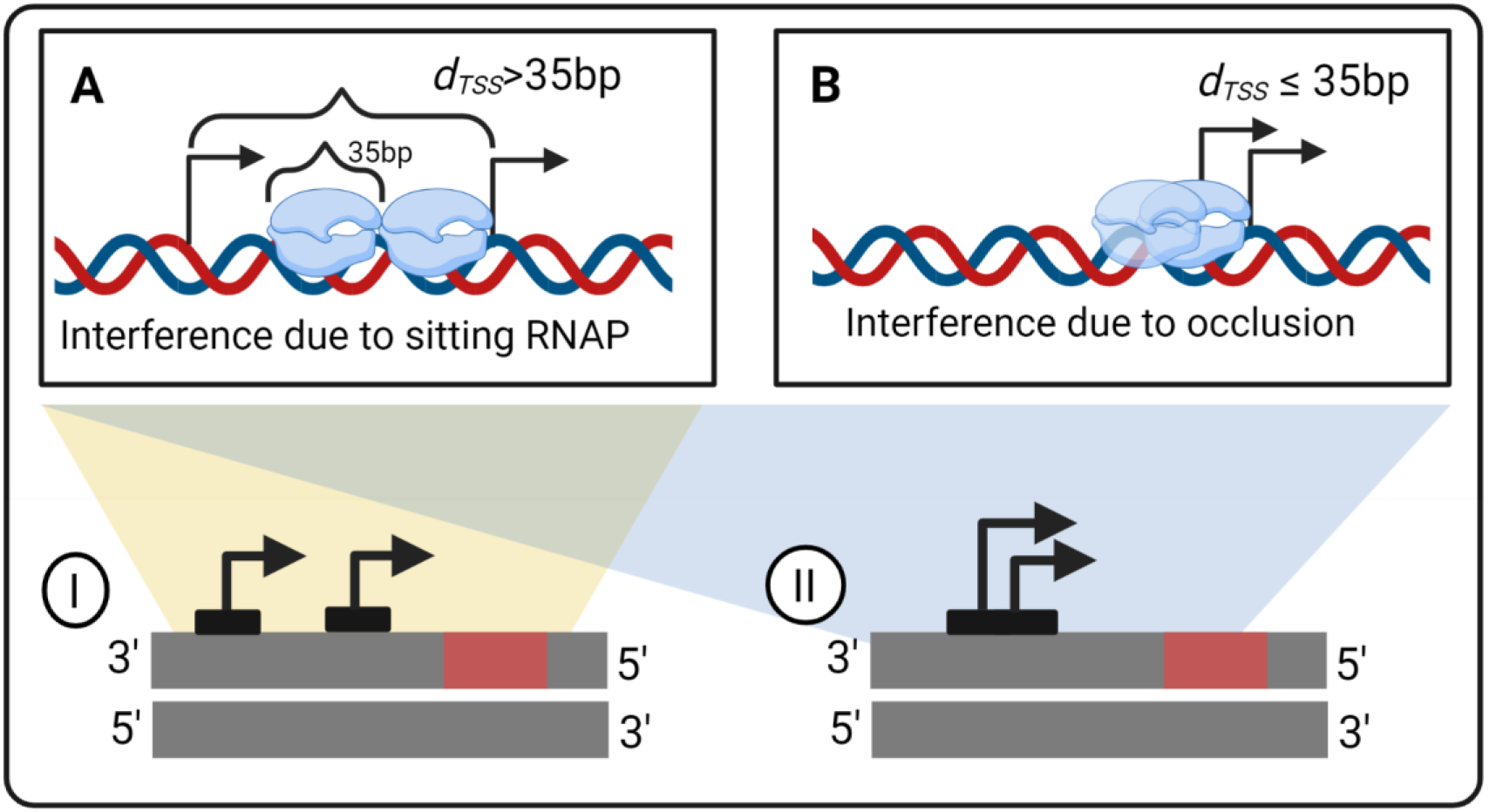
Interference between tandem promoters with different arrangements relative to each other and to neighbour genes. **(A)** Interference by an RNAP occupying the downstream promoter on the activity of the elongating RNAP from upstream promoter. The TSSs need to be at least 36 bp apart (the length occupied by an RNAP when in OC, [23, 25]) **(B)** Interference by occlusion of one of the promoter’s TSS by an RNAP on the TSS of the other promoter. The distance between the TSSs need to be ≤ 35 bp apart. Blue clouds are RNAPs. Black arrows sit on TSSs and point towards the direction of transcription elongation. Arrangements **(I-II)** of two promoters studied in the manuscript in tandem formation are represented. The red rectangles are the protein coding regions. We studied only the natural tandem promoters that neither overlap with nor have in between another gene (arrangements I and II, which differ based on whether the promoter regions overlap or not). Other arrangements (not considered in this study) are shown in Fig. S1 in the S2 Appendix. Figure created with BioRender.com.

The numbers of these arrangements in *E. coli* are shown in Table S8 in the S3 Appendix. From the measurements of these genes’ protein levels, we then establish a model that we use to explore the state space of potential dynamics under the control of tandem promoters (Fig 2 illustrates our workflow).

**Fig 2.**
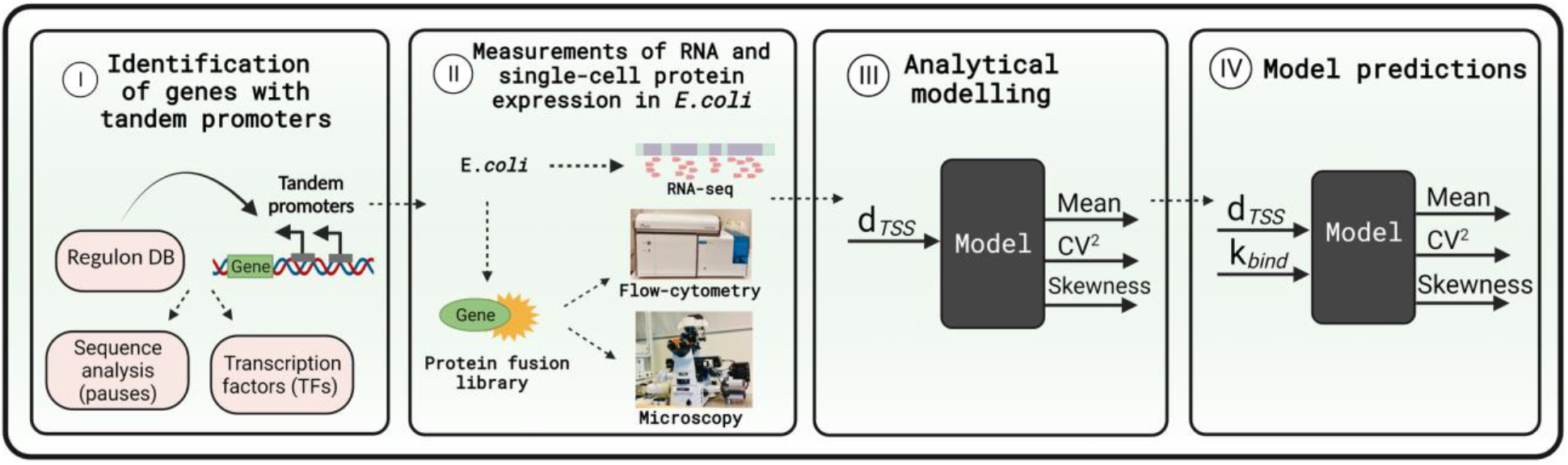
Workflow. (I) We identified genes controlled by tandem promoters in Regulon DB. (II) Next, we measured the single-cell protein levels of those genes with arrangements I and II that are tagged in the YFP strain library [28]. We also measured the mean RNA fold changes of these genes over time (S1 Appendix, section ‘RNA-seq measurements and data analysis’). (III) We used the single-cell data to tune the model. (IV) Finally, we used the model to explore the state space of protein expression. Figure created with BioRender.com.

## Results

*E. coli* has 831 genes controlled by two or more promoters in tandem formation (RegulonDB and section ‘Selection of natural genes controlled by tandem promoters for flow-cytometry’ in the S1 Appendix). However, to study the dynamics of genes controlled by tandem promoters, we focused on only 102 of them, because their activity is expected to be undisturbed by neighboring genes in the DNA (arrangements I and II in Fig 1), for reasons described in section ‘Selection of natural genes controlled by tandem promoters for flow-cytometry’ in the S1 Appendix.

Further, these promoters do not have specific short nucleotide sequences capable of affecting RNAP elongation (section ‘Pause sequences’ in the S4 Appendix). Also, the 102 genes expressed by these promoters are not overrepresented in a particular biological process (section ‘Over-representation test’ in the S4 Appendix). From time-lapse RNA-seq data (S1 Appendix, section ‘RNA-seq measurements and data analysis’), we also did not find evidence that their dynamics are affected by their input transcription factors (TFs) in our measurement conditions (section ‘Input-output transcription factor relationships’ in the S4 Appendix) nor by H-NS in a consistent manner (section ‘Regulation by H-NS’ in the S4 Appendix). Finally, they do not exhibit any particular TF network features (Table S3 in the S3 Appendix). As such, neither input TFs nor specific nucleotide sequences are considered in the model below. In addition to all of the above, we found no correlations between the shortest distance from the TSS of upstream promoters from the oriC region in the DNA and expression levels (section ‘Relationship with the oriC region’ in the S4 Appendix).

### Model of gene expression controlled by tandem promoters

RNAPs bind, slide along, and unbind from a promoter several times until, eventually, one of them finds the TSS [29–30], commits to OC at the TSS, and initiates transcription elongation.

Reactions (1a1) are a 4-step (I-IV) model of transcription [20, 31]. The forward reaction in step I in (1a1) models RNAP binding to a free promoter (*P_free_*), which becomes no longer free albeit the RNAP might not yet have reached the TSS. This state, pre-finding of the TSS, is here named *P_bound_* and its occurrence increases with RNAP concentration, *[R]*. Next, as it percolates the DNA, the RNAP should find and stop at the nearest TSS and form a closed complex (CC) with the DNA (step II, Reaction 1a1). CCs are unstable, i.e. reversible [22] (reaction 1a2) but, eventually, one of them will commit to OC irreversibly [32], via step III, Reaction 1a1 [21–22]. It follows RNAP escape from the TSS, freeing the promoter (step IV, Reaction 1a1) [33–37]. Then, the RNAP elongates (*R_elong_*) until producing a complete RNA (reaction 1a3) and freeing itself.

These set of reactions usually model well stochastic transcription dynamics [20]. However, if two promoters are closely spaced in tandem formation, they can interfere [38]. Figure 3 shows sequences of events that can lead to interference between tandem promoters, not accounted for by the model above.

**Fig 3.**
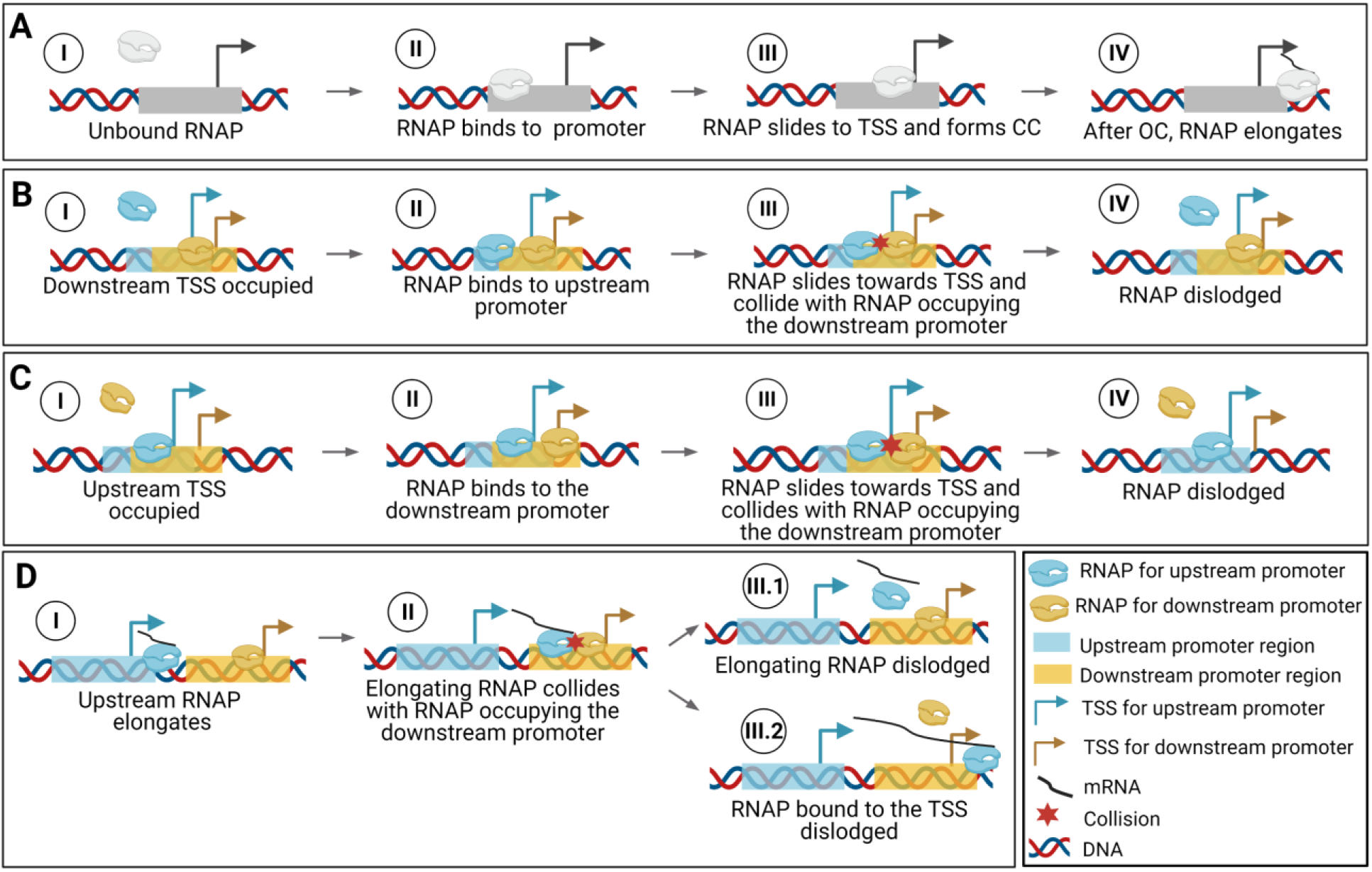
Events leading to transcriptional interference between tandem promoters. (**A**) Sequence of events in transcription in isolated promoters. A similar set of events occurs in tandem promoters, if only one RNAP interacts with them at any given time. (**B / C**) Interference due to the occlusion of the *downstream* / *upstream* promoter by a bound RNAP, which will impede the incoming RNAP from binding to the TSS. (**D**) Interference of the activity of the RNAP incoming from the upstream promoter by the RNAP occupying the downstream promoter. One of these RNAPs will be dislodged by the collision. Created with BioRender.com.

From Figure 3, if the TSSs are sufficiently close, the occupancy of one TSS by an RNAP will occlude the other TSS, blocking its kinetics [18]. This is accounted for by reaction 1a5, which competes with CC formation in reaction 1a1. Its rate constant, k_occlusion_, is defined in the next section. In (1a5), ‘u/d’ stands for occlusion of the upstream promoter by an RNAP on the TSS of the downstream promoter.

Instead, if the TSSs are not sufficiently close, they will still interfere since the elongating RNAP (*R_elong_*) starting from the upstream promoter can collide with RNAPs on the TSS of the downstream promoter. This can dislodge either RNAP via (reaction 1a4) or (reaction 2a3), depending on the sequence-dependent binding strength of the RNAP to the TSS [9].

Finally, once reaction 1a1 occurs, either reaction 1a3 or 1a4 occur. To tune their competition, we introduced the terms ω_d_ and (1-ω_d_) in their rate constants, with ω_d_ being the fraction of times that an elongating RNAP from an upstream promoter finds an RNAP occupying the downstream promoter. Meanwhile, ‘*f*’ is the fraction of times that the RNAP occupying the downstream promoter falls-off due to the collision with an elongating RNAP, whereas ‘*1-f’* is the fraction of times that it is the elongating RNAP that falls-off.

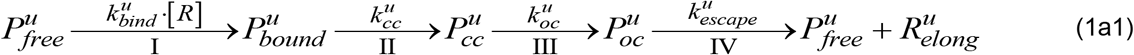

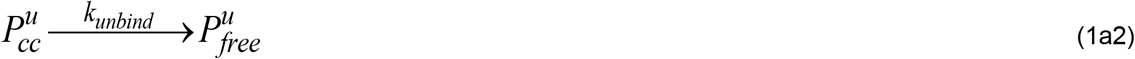

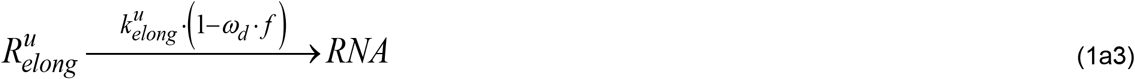

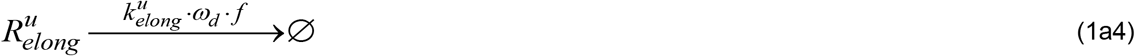

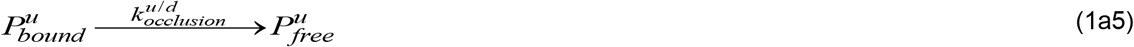

Next, we reduced the model and derived its analytical solution. First, since *P_cc_* completion is expected to be faster than *P_bound_* completion ([10] and references within) we merged them into a single state, *P_occupied_*, which represents a promoter occupied by an RNAP prior to commitment to OC, whose time length is similar to *P_bound_*.

Similarly, in standard growth conditions, the occurrence of multiple failures in escaping the promoter [46] per OC completion should only occur in promoters with the highest binding affinity to RNAP. Thus, in general promoter escape should be faster than OC [20, 32]. We thus merged OC and promoter escape into one step named ‘events *after* commitment to OC’, with a rate constant *kafter*. The simplified model is thus:

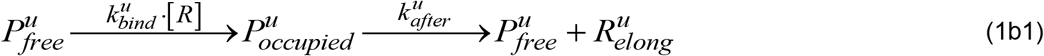

These two steps are not merged since only the first differs with RNAP concentration [20, 26,39]. Further, reports [40–41] indicate that *E. coli* has ~100-1000 RNAPs free for binding at any moment but ~4000 genes, suggesting that the number of free RNAPs is a limiting factor.

Finally, we merge (1a2), (1a5) and (1b1) in one multistep without affecting the model kinetics:

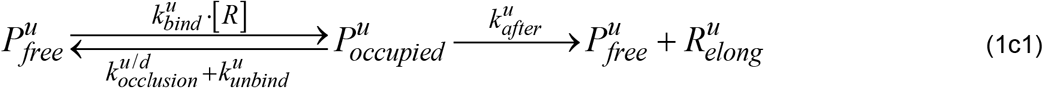

Overall, this reduced model of transcription of upstream promoters has a multistep reaction of transcription initiation (1c1), a reaction of transcription elongation (1a3) and a reaction for failed elongation due to RNAPs occupying the downstream promoter (1a4).

Regarding RNA production from the downstream promoter, it should either be affected by occlusion if *d_TSS_* ≤ 35, or by RNAPs elongating from the upstream promoter if *d_TSS_* > 35 (Fig 3). We thus use reactions (2a1), (2a2), and (2a3) to model these promoters’ kinetics:

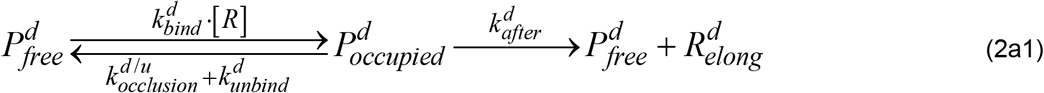

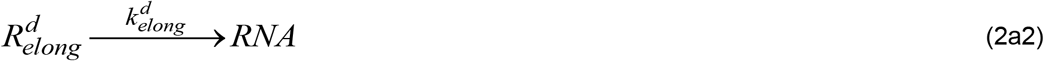

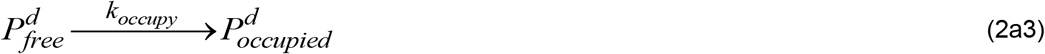

Finally, one needs to include a reaction for translation (reaction 3), as a first order process since protein numbers follow RNA numbers linearly (Fig S6 in the S2 Appendix), and reactions for RNA and protein decay accounting for degradation and for dilution due to cell division (reactions 4a and 4b, respectively). TF regulation is not included as noted above (Figs S3 and S4A in the S2 Appendix).

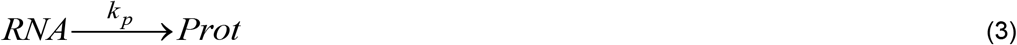

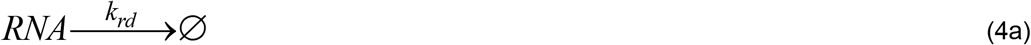

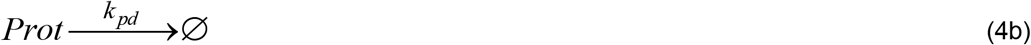

### Transcription interference by occlusion

In a pair of tandem promoters, the *k_occlusion_* of one of them should increase with the fraction of time that the other one is occupied. Further, it should decrease with increasing *d_TSS_* between the two promoters’ TSS. We thus define *k_occlusion_* for the upstream (Eq. 5a) and downstream (Eq. 5b) promoters, respectively as:

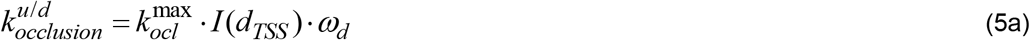

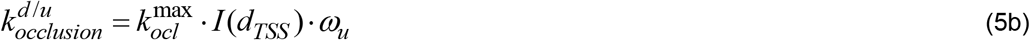

Here, 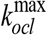 is the maximum occlusion possible. It occurs when the two TSSs completely overlap each other (*d_TSS_ = 0*) *and* the TSS of the ‘other’ promoter is always occupied. Meanwhile, *I(d_TSS_)* models distance-dependent interference.

We tested four models of interference: ‘exponential 1’, ‘exponential 2’, ‘step’, and ‘zero order’ (Table 1). The first two assume that the effects of occlusion decrease exponentially with *d_TSS_* (first and second order dependency, respectively).

**Table 1.** Potential models of transcriptional interference due to promoter occlusion considered.

| Interference by occlusion | $I(d_{TSS})$ | $k_{occlusion}$ |
| --- | --- | --- |
| Exponential 1 (“Exp1”) | $e^{-(b_1 \cdot d_{TSS})}$ | $k_{ocl}^{\max} \cdot e^{-(b_1 \cdot d_{TSS})} \cdot \omega$ |
| Exponential 2 (“Exp2”) | $e^{-(b_1 \cdot d_{TSS} + b_2 \cdot d_{TSS}^2)}$ | $k_{ocl}^{\max} \cdot e^{-(b_1 \cdot d_{TSS} + b_2 \cdot d_{TSS}^2)} \cdot \omega$ |
| Step (“Step”) | $1 - \frac{1}{1 + e^{-m \cdot (d_{TSS} - L)}}$ | $k_{ocl}^{\max} \cdot \left(1 - \frac{1}{1 + e^{-(d_{TSS} - L)}}\right) \cdot \omega$, for $m = 1 \text{ bp}^{-1}$ |
| Zero order ("ZeroO") | $k$ | $k_{ocl}^{\max} \cdot \omega$ |

Meanwhile, the ‘Step’ model assumes that interference only occurs precisely in the region in the DNA occupied by the RNAP when in OC formation. For this, it uses a logistic equation to build a continuous step function, where *L* is the length of DNA (in bp) occupied by the RNAP in OC. As such, L tunes at what *d_TSS_* the step occurs, while *m* is the steepness of that step (set to 1 bp^−1^).

Finally, the ‘Zero order’ model assumes (unrealistically) that interference by occlusion, is independent of *d_TSS_*. Fig S7 in the S2 Appendix shows how *k_occlusion_* differs with *d_TSS_* in each model, for various parameter values.

Finally, ω is the fraction of time that the ‘other’ promoter is occupied. It ranges from 0 (no occupancy) to 1 (always occupied). It is estimated for upstream and downstream promoters as:

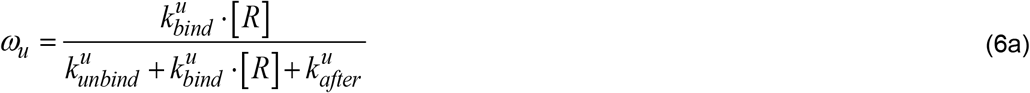

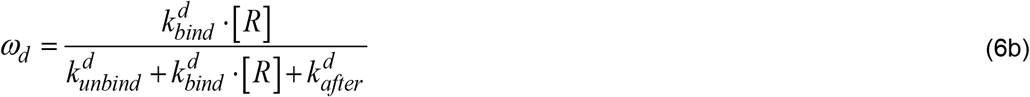

Similarly, 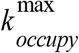 if is the maximum possible interference due to RNAPs occupying the downstream promoter, *k_occupy_* is defined as:

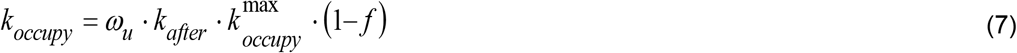

### Analytical solution of the moments of the single-cell protein numbers

Next, we derived an analytical solution of the expected mean single-cell protein numbers at steady state, *M_P_*, which is later tuned to fit the empirical data. For any gene, regardless of the underlying kinetics of transcription, *k_r_* is the *effective* rate of RNA production. Based on the reactions above, the mean protein numbers in steady state will be (see sections “Analytical model of mean RNA levels controlled by a single promoter in the absence of a closely spaced promoter” and “Derivation of mean protein numbers at steady state produced by a pair of tandem promoters” in the S1 Appendix):

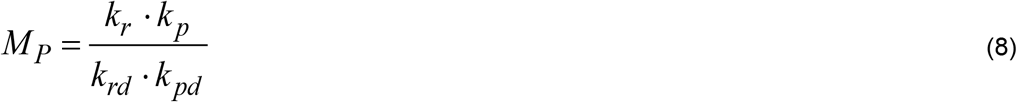

This equation applies to a pair of tandem promoters as well. In that case, assuming that *k_bind_* of the two tandem promoters is similar, we have:

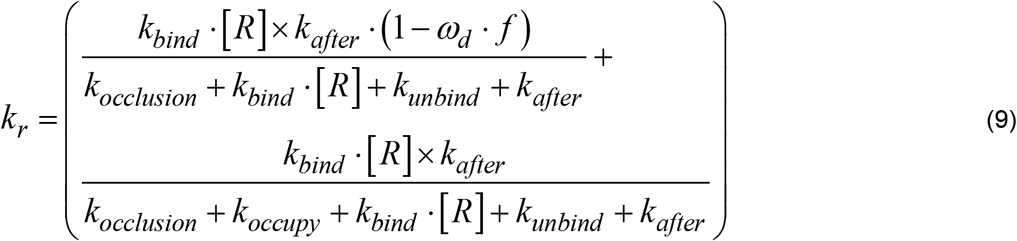

To derive the other moments, we considered that empirical single-cell protein numbers in *E. coli* are well fit by negative binomials [28]. Consequently, *M_p_* and the squared coefficient of variation 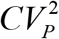, should be related as (Equations S28 to S38 in the S1 Appendix):

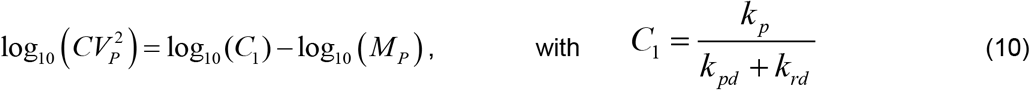

This relationship matches empirical data at the genome wide level, except for genes with high transcription rates [54]. Additionally, we further derived a relationship (Section ‘CV^2^ and Skewness of single-cell protein expression of a tandem promoter’s model’ in the S1 Appendix) between *M_P_* and the skewness, *S_P_*, of the single-cell distribution of protein numbers:

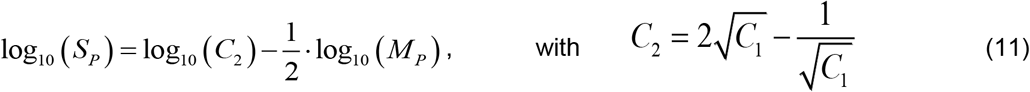

### Single-cell distributions of protein numbers

To validate the model, we measured by flow-cytometry the single-cell distributions of protein fluorescence of 30 out of the 102 genes known to be controlled by tandem promoters (with arrangements I and II). Measurements were made in 1X and 0.5X media (3 replicates per condition) using cells from the YFP strain library (section ‘Strains and Growth Conditions’ in the S1 Appendix). Data from past studies show that, in these 30 genes, RNA and protein numbers are well correlated (Fig S6 in the S2 Appendix) in standard growth conditions. Past studies also suggest that most of these genes are active during exponential growth (~95% of our 30 genes selected should be active, according to data in [42] using SEnd-seq technology).

Single-cell distributions of protein expression levels are shown in Fig 4A for one of these genes as an example. The raw data from all 30 genes (only one replicate) are shown in Fig S8 in the S2 Appendix. Finally, the mean, CV^2^ and skewness for each gene, obtained from the triplicates, are shown in Excel sheets 1 and 2 in the S6 Table. In addition, we also show this mean, CV^2^ and skewness after subtracting the first, second, and third moments of the single-cell distribution of the fluorescence of control cells, which do not express YFP (Sheets 3, 4 in the S6 Table) (Section ‘Subtraction of background fluorescence from the total protein fluorescence’ in the S1 Appendix).

**Fig 4.**
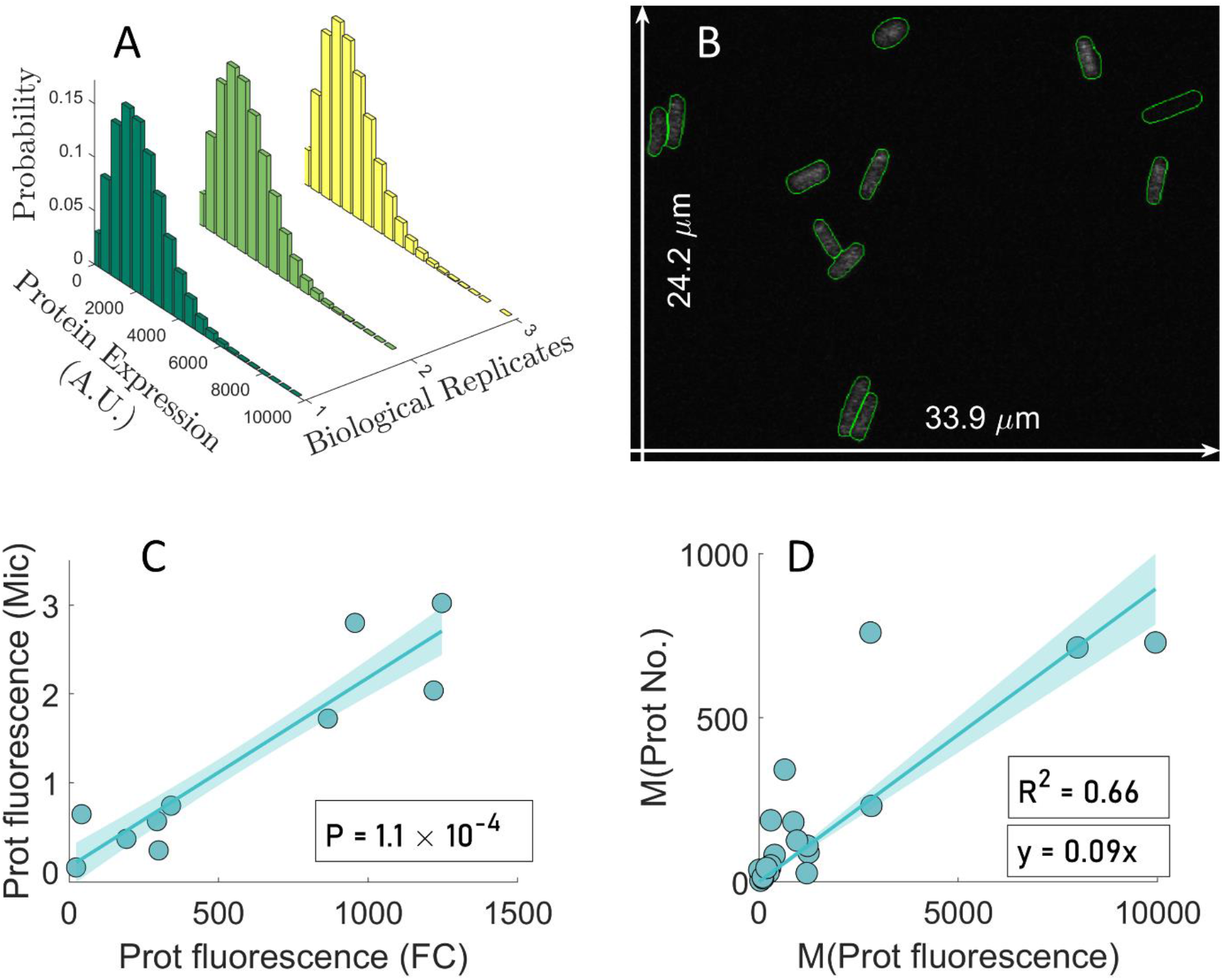
Single cell protein numbers by microscopy and flow-cytometry. (A) Example single-cell distributions (3 biological replicates) of fluorescence (in arbitrary units) of cells with a YFP tagged gene controlled by a pair of tandem promoters obtained by flow-cytometry, ‘FC’. (B) Example confocal microscopy image of cells overlapped by the results of cell segmentation from the corresponding phase contrast image. The two white arrows show the dimensions of the image, for scaling purposes. (C) Mean single-cell protein fluorescence of 10 genes (Table S7 in the S3 Appendix) when obtained by FC plotted against when obtained by microscopy, ‘Mic’. (D) Mean single-cell protein fluorescence (own measurements) plotted against the corresponding mean single-cell protein numbers reported in [28]. From the equation of the best fitting line without y-intercept (y-intercept = 0), we obtained a scaling factor, *sf,* equal to 0.09.

Based on the analysis of the data of these 30 genes, we removed from subsequent analysis those genes (5 in 1X and 14 in 0.5X) whose mean, variance, or third moment of their protein fluorescence distributions are lower than in control cells (not expressing YFP), i.e., than cellular autofluorescence (Sheets 3, 4 in S6 Table). As such, only one gene studied here (in condition 1X alone) codes for a protein that is associated to membrane-related processes, which might affect its quantification (section ‘Proteins with membrane-related positionings’ in S4 Appendix). As such, we do not expect this phenomenon to influence our results significantly. The data from these genes removed from further analysis is shown in Fig S6 in S2 Appendix alone, for illustrative purposes.

We started by testing the accuracy of the background-subtracted flow-cytometry data by confronting it with microscopy data (also after background subtraction, see section ‘Microscopy and Image Analysis’ in the S1 Appendix). We collected microscopy data on 10 out of the 30 genes (Table S7 in the S3 Appendix). The microscopy measurements of the mean single-cell fluorescence expressed by these genes (example image in Fig. 4B), were consistent, statistically, with the corresponding data obtained by flow-cytometry (Fig 4C).

Next, we converted the fluorescence distributions from flow-cytometry (25 genes in 1X and 16 genes in 0.5X) into protein number distributions. In Fig 4D we plotted our measurements of mean protein fluorescence in 1X against the protein numbers reported in [28] for the same genes, in order to obtain a scaling factor (sf = 0.09). Using sf, we estimated *M_P_*, 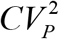, and *S_P_* of the distribution of protein numbers expressed by the tandem promoters in (Sheets 5, 6 in S6 Table) (Section ‘Conversion of protein fluorescence to protein numbers’ in S1 Appendix).

To test the robustness of the estimation of the scaling factor, we also estimated a scaling factor from 10 other genes present in the YFP strain library [28] (listed in Table S2 in S3 Appendix). These genes were selected as described in the section ‘Selection of natural genes controlled by single promoters’ in S1 Appendix. Using the data from this new gene cohort (Supplementary Figure S9A in S2 Appendix) reported in S7 Table, we estimated a scaling factor of 0.08, supporting the previous result. Meanwhile, since when merging the data from tandem and single promoters, the resulting scaling factor equals 0.09 (Supplementary Figure S9B in S2 Appendix), we opted for using 0.09 from here onwards.

We also tested how sensitive the estimated scaling factor is to the removal of data points. Specifically, for 1000 times, we discarded N randomly selected data points, and estimated the resulting scaling factor. We then compared, for each N, the mean and the median of the distribution of 1000 scaling factors (Supplementary Figure S10 in S2 Appendix). Since the median is not sensitive to outliers, if mean and median are similar, one can conclude that the scaling factor is not biased by a few data points. Visibly, the mean and the median only start differing for N larger than 6, which corresponds to nearly 30% of the data.

### Log-log relationship between the mean single-cell protein numbers of tandem promoters and the other moments

We plotted *M_P_* against 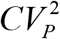 and *S_P_* in log-log plots, in search for the fitting parameters, ‘*C_1_*’ and ‘*C_2_’*, to estimate the rate of protein production per RNA (equation 10). To increase the state space covered by our measurements, in addition to M9 media (named ‘1X’), we also used diluted M9 media (named ‘0.5X’), known to cause cells to have lower RNAP concentrations (Fig. 5A) (Section ‘Strains and growth conditions’ in the S1 Appendix), without altering the division rate (Figs. S11A and S11B in the S2 Appendix). We note that 1X and 0.5X only refer to the degree of dilution of the original media and not to how much RNAP concentration and consequently, protein concentrations, were reduced by media dilution. From the same figures, we attempted stronger dilutions, but no further decreases in RNAP concentration were observed and the growth rate decreased.

**Fig 5.**
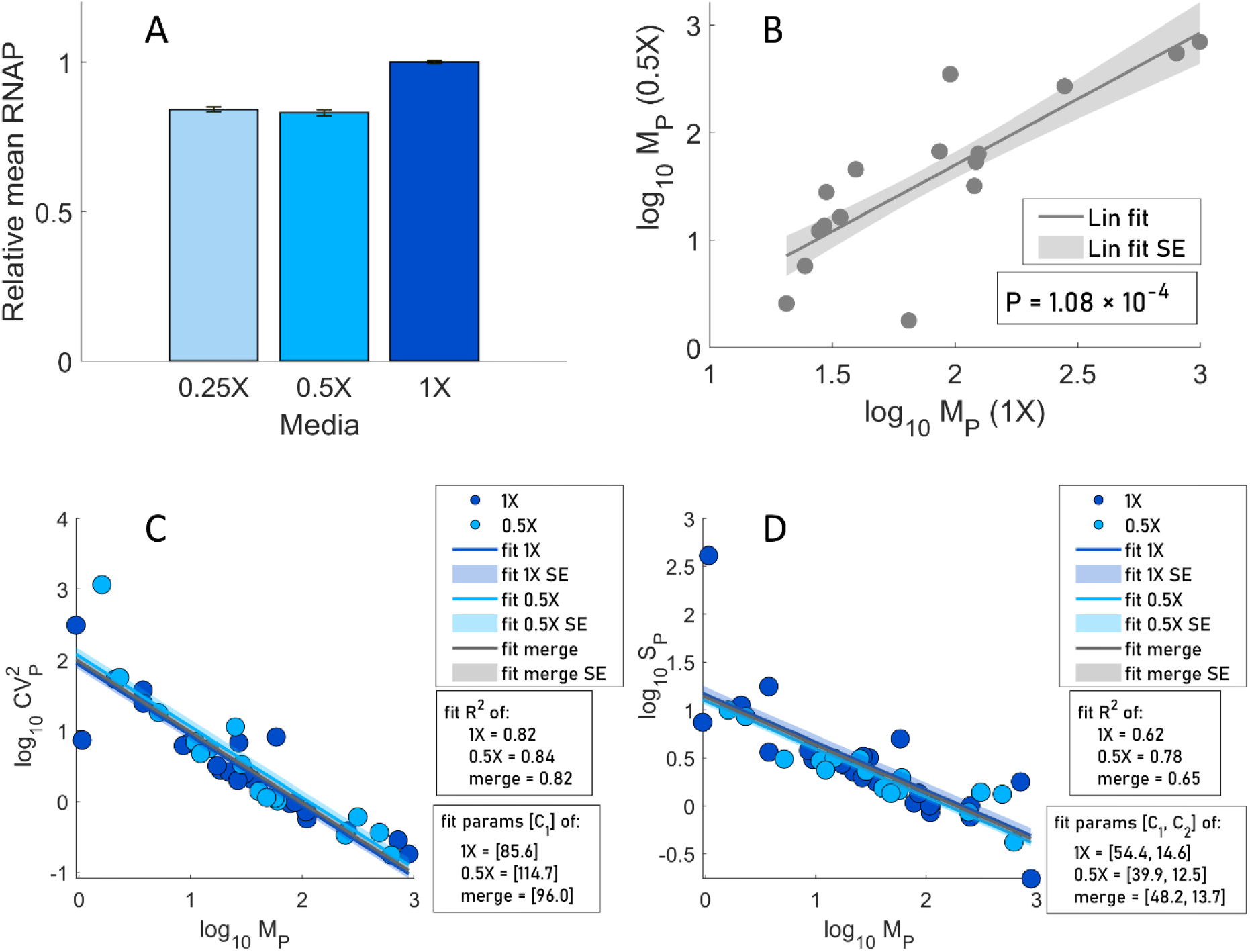
Relative RNAP concentrations along with the relationships between the moments of the single cell distributions of protein numbers. (A) Relative RNAP levels measured by flow-cytometry (Section ‘flow-cytometry’ in the S1 Appendix) in three media. (B) Scatter plot between MP in M9 (1X) and diluted M9 (0.5X) media. Also shown are the best fitting line and standard error and p-value for the null hypothesis that the slope is zero. (C) *M_P_* vs 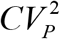 and (D) *M_P_* vs *S_P_* of single-cell protein numbers of genes with tandem promoters in M9 (1X) and M9 diluted (0.5X) media. The lines and their shades are the best fitting lines and standard errors, respectively. ‘Merge’ stands for data from both 0.5X and 1X conditions.

Next, from Fig 5B, most genes (of those expressing tangibly in both media) suffered similar reductions (well fit by a line) in protein numbers with the media dilution, as expected by the model of gene expression (Equations 8 and 9). This linear relationship could also be interpreted as evidence that the difference in expression of these genes between the two conditions is not affected by TFs in our measurement conditions. Namely, if TF influences existed, and TF numbers changed, they would likely be diversely affected by their output genes (weakly and strongly activated, repressed, etc.) and, thus, our proteins of interest would not have changed in such similar manners (linearly).

Meanwhile, as in [44–45], 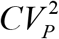 decreases linearly with *M_P_* (log-log scale), irrespective of media (R^2^ > 0.8 in all fitted lines), in agreement with the model (Fig 5C). Fitting Equation 10 to the data, we extracted *C_1_* in each condition. *S_P_* also decreases linearly with *M_P_*, irrespective of the media (Fig 5D). Similar to above, Equation 11 was fitted to each data set and *C_1_* and *C_2_* were obtained (R^2^ > 0.6 for all lines).

Since *C_1_* from Fig 5C and 5D differed slightly (likely due to noise), we instead obtained *C_1_* and *C_2_* values that maximized the mean R^2^ of both plots. Using ‘fminsearch’ function in MATLAB [46], we obtained *C_1_* = 72.71 and *C_2_* = 16.94 (R^2^ of 0.80 and 0.61, respectively) for Fig 5C and Fig 5D, respectively.

### Inference of parameter values and model predictions as a function of *d_TSS_*

We next used the model, after fitting, to predict how *d_TSS_* and the promoters’ occupancy regulate the moments of the single-cell distribution of protein numbers (*M_P_*, 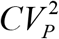, and *S_P_*) under the control of tandem promoters. We started by assuming the parameter values from the literature listed in Table 2 and tuned the remaining parameters.

**Table 2.** Parameter values imposed identically on all models.

| Parameter description | Parameter | Value | References |
| --- | --- | --- | --- |
| Inverse of the mean time to complete OC | $k_{after}$ | $0.005 \text{ s}^{-1}$ | Differs between promoters. Since empirical data lacks, we used the data from *in vivo* single RNA measures for Lac-Ara-1 [20]. |
| RNA and protein dilution due to division | $k_{dil} = \frac{\ln(2)}{D}$ | $1.005 \times 10^{-4} \text{ s}^{-1}$ | Legend of Fig S8 |
| RNA degradation | $k_{rdeg}$ | $2.3 \times 10^{-3} \text{ s}^{-1}$ | [28] |
| RNA decay due to dilution from cell division and due to degradation | $k_{rd} = k_{rdeg} + k_{dil}$ | $2.4 \times 10^{-3} \text{ s}^{-1}$ | From row 2. |
| Protein degradation | $k_{pdeg}$ | $2.93 \times 10^{-5} \text{ s}^{-1}$ | [47], estimates it to be from $\sim 6 \times 10^{-5}$ to $\sim 2 \times 10^{-5}$. We used the value in [48], in that interval. |
| Protein decay due to dilution by cell division and degradation | $k_{pd} = k_{pdeg} + k_{dil}$ | $1.3 \times 10^{-4} \text{ s}^{-1}$ | From rows 2 and 5. |
| Fall-off probability of the RNAP occupying the downstream promoter | $f$ | 50% (0.5) | Set here (likely sequence-dependent) |
| Protein production rate constant | $k_p = C_1 \times (k_{pd} + k_{rd})$ | $0.18 \text{ s}^{-1}$ | $C_1$ is estimated here. |
| Free RNAP per cell | $[R]$ | 144/cell in 1X and 120/cell in 0.5X media | See main text. |

To set the RNAP numbers in Table 2, we considered that the RNAPs affecting transcription rates are the *free* RNAPs in the cell, and that, for doubling times of 30 min in rich medium, there are ~1000 free RNAPs per cell [41]. Meanwhile, for doubling times of 60 min in minimal medium, there are ~144 [40]. In both our media, we observed a doubling time of ~115 mins (Fig. 5B). Thus, we expect the free RNAP in 1X to also be ~144/cell or lower. Meanwhile, in 0.5X, we measured the RNAP concentration to be 17% lower than in 1X (Fig. 5A) and no morphological changes. Thus, we assume the free RNAP in 0.5X to equal ~120/cell.

Next, we fitted the equations (8) and (9) relating *d_TSS_* with log_10_ (*MP*) in all interference models (Table 1), using the data on *MP* in 1X medium (Fig 6A) and the ‘fit’ function of MATLAB. For this, we set 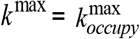 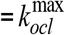, for simplicity, as well as realistic bounds for each parameter to infer. To avoid local minima, we performed 200 searches, each starting from a random initial point, and selected the one that maximized *R^2^*. Results are shown in Table 3.

**Table 3.** Parameter values inferred for each model.

| Interference model | Inferred parameter values | Average R<sup>2</sup> (M, CV<sup>2</sup>, S) 1X medium | Average R<sup>2</sup> (M, CV<sup>2</sup>, S) 0.5X medium |
| --- | --- | --- | --- |
| Exponential 1 | $k_{bind} \cdot [R] = 1.09 \times 10^{-2} \text{ s}^{-1} \times (\text{cell vol})^{-1}$<br>$k_{bind} = 7.53 \times 10^{-5} \text{ s}^{-1}$ | 0.21 (Figs. 6A, 6B, and 6C) | 0.09 (Figs. 6D, 6E, and 6F) |
| | $k_{unbind} = 0.84 \text{ s}^{-1}$<br>$k^{\max} = 677.7 \text{ s}^{-1}$<br>$b_1 = 5.08 \times 10^{-2} \text{ bp}^{-1}$ | | |
| Exponential 2 | $k_{bind} \cdot [R] = 9.71 \times 10^{-3} \text{ s}^{-1} \times (\text{cell vol})^{-1}$<br>$k_{bind} = 6.74 \times 10^{-5} \text{ s}^{-1}$<br>$k_{unbind} = 0.80 \text{ s}^{-1}$<br>$k^{\max} = 554.8 \text{ s}^{-1}$<br>$b_1 = 7.92 \times 10^{-8} \text{ bp}^{-1}$<br>$b_2 = 1.47 \times 10^{-3} \text{ bp}^{-2}$ | 0.25 (Figs. 6A, 6B, and 6C) | 0.12 (Figs. 6D, 6E, and 6F) |
| Step | $k_{bind} \cdot [R] = 6.62 \times 10^{-3} \text{ s}^{-1} \times (\text{cell vol})^{-1}$<br>$k_{bind} = 4.60 \times 10^{-5} \text{ s}^{-1}$<br>$k_{unbind} = 0.49 \text{ s}^{-1}$<br>$k^{\max} = 313.4 \text{ s}^{-1}$<br>$L = 35.11 \text{ bp}$ (by best fitting, which corresponds to 35 bp) | 0.35 (Figs. 6A, 6B, and 6C) | 0.15 (Figs. 6D, 6E, and 6F) |
| zero order | $k_{bind} \cdot [R] = 4.63 \times 10^{-3} \text{ s}^{-1} \times (\text{cell vol})^{-1}$<br>$k_{bind} = 3.22 \times 10^{-5} \text{ s}^{-1}$<br>$k_{unbind} = 0.57 \text{ s}^{-1}$<br>$k^{\max} = 6.48 \text{ s}^{-1}$ | -0.007 (Figs. 6A, 6B, and 6C) | -0.12 (Figs. 6D, 6E, and 6F) |

**Fig 6.**
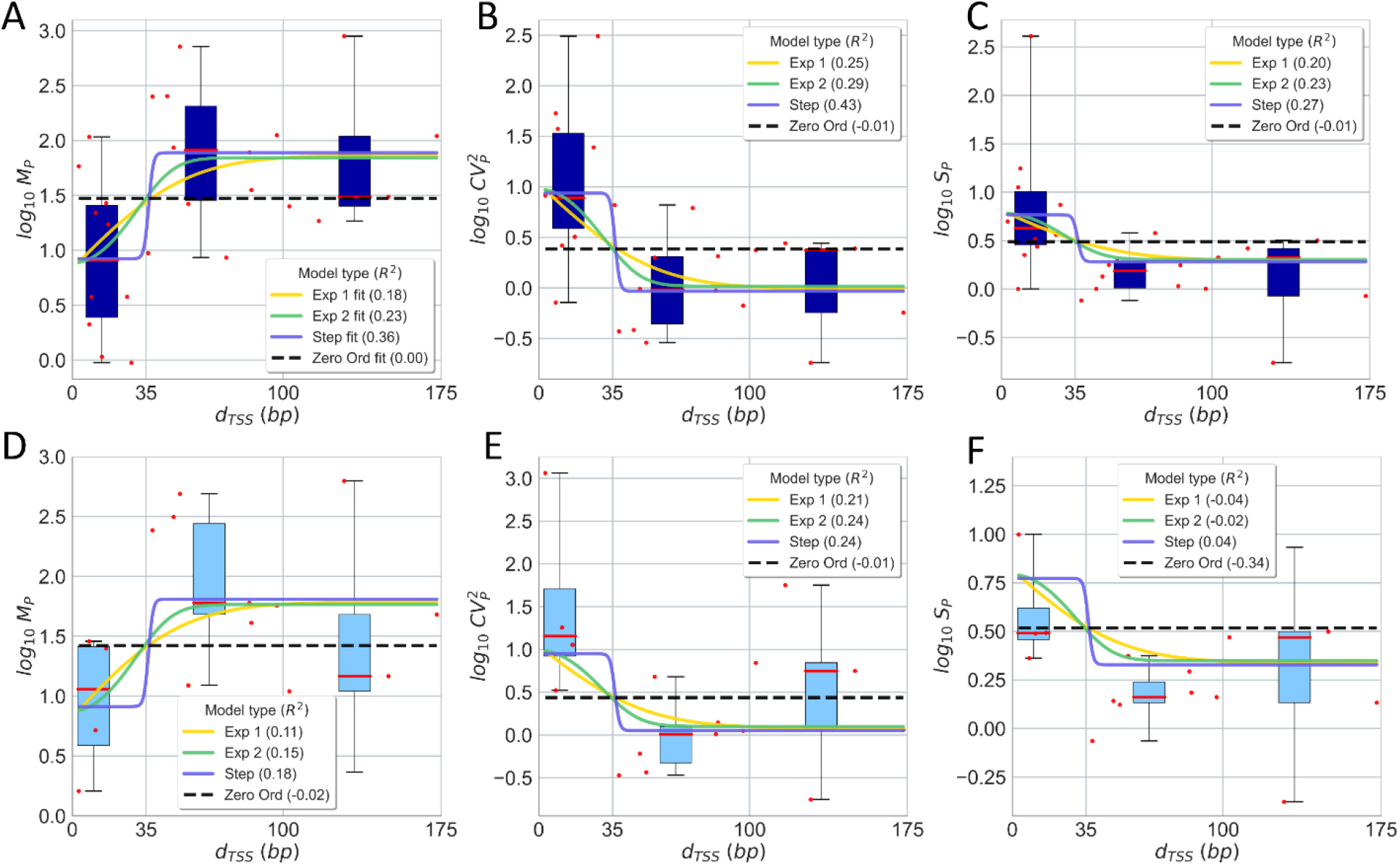
Empirical data and analytical model of how *d_TSS_* influences the single-cell protein numbers of genes controlled by tandem promoters. **(A)** Mean, **(B)** *CV^2^*, and **(C)** *S* of single protein numbers in the **1X** media as a function of *d_TSS_*. **(D)**, **(E),** and **(F)** show the same for the **0.5X** media, respectively. Each red dot is the mean from 3 biological repeats for a pair of promoters (S6 Table). The dots were also grouped in 3 ‘boxes’ based on their *d_TSS_*. In each box, the red line is the median and the top and bottom are the 3^rd^ and 1^st^ quartiles, respectively. The vertical black bars are the range between minimum and maximum of the red dots. In **A**, all lines are best fits. In **B**, **C**, **D**, **E**, and **F**, all lines are model predictions, based on the parameters used to best fit **A**. The insets show the *R^2^* for each model fit and prediction.

Next, we inserted all parameter values (empirical and inferred) in Equations (10) and (11) to predict 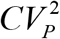 and *S_P_* in 1X medium (Figs 6B and 6C). Also, we inserted the same parameter values and the estimated RNAP numbers in 0.5X medium in equations (8–11) to obtain the analytical solutions for *M_P_*, 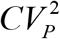 and *S_P_* for 0.5X medium (Figs 6D, 6E and 6F).

From Fig. 6, the data is ‘noisy’, which suggests that it is not possible to establish if the models are significantly different. As such, here we only select the one that best explains the data, based on the R^2^ values of the fittings. Table 3 shows the mean *R^2^* for *M_P_*, 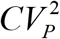, and *S_p_* when confronting the model with the data. Overall, from the R^2^ values, the step model is the one that best fits the data. Meanwhile, the ‘ZeroO’ model is the least accurate, which supports the existence of distinct kinetics when *d_TSS_* is smaller or larger than 35 nucleotides, which is the length of the RNAP when committed to OC on the TSS [23–25].

In summary, the proposed model of expression of genes under the control of a pair of tandem promoters is based on a standard model of transcription of each promoter, which are subject to interference, either due to occlusion of the TSSs or by RNAP occupying the downstream promoter on the TSS of the downstream promoter. The influence of each occurrence of these events is well modeled by linear functions of TSS occupancy times, while their dependency on *d_TSS_* is modeled by a continuous step function. If *d_TSS_* is larger than 35 bp, effects from the RNAP occupying the downstream promoter can occur, else occlusion can occur.

We then confronted the analytical solutions of the step model with stochastic simulations (Section ‘Stochastic simulations for the step inference model’ in the S1 Appendix). We first assumed various *d_TSS_*, but fixed *k_bind_*, for simplicity. Visibly, *M_P_*, 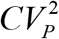, and *S_p_* of the stochastic simulations are well-fitted by the analytical solution, supporting the initial assumption that 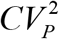, and *S_p_* follow a negative binomial (Fig S13 in the S2 Appendix).

However, natural promoters are expected to differ in *k_bind_* as they differ in sequence [49–50]. Thus, we introduced this variability and studied whether the analytical model holds. To change the variability, we obtained each *k_bind_* from gamma distributions (means shown in Table 3 and CVs in Table S9 in the S3 Appendix). We chose a gamma distribution since its values are non-negative and non-integer (such as rate constants). Meanwhile, all parameters of the step model, aside from *k_bind_*, are obtained from Tables 2 and 3. For *d_TSS_* ≤ 35 and *d_TSS_* > 35, and each CV considered, we sampled 10.000 pairs of values of *k_bind_* · [*R*], and calculated M, CV^2^ and S for each of them. Next, we estimated the average and standard deviation of each statistics. From Fig S14 in the S2 Appendix, if *CV* (*k_bind_*) < 1, the analytical solution is robust. In that the standard error of the mean is smaller than M_P_/3. Notably, for such C*V*, the strength of the two paired promoters would have to differ unrealistically by more than 2000%, on average (Table S9 in the S3 Appendix). Thus, we find the analytical solution to be reliable.

From our estimation of *k_p_*, we further estimated a protein-to-RNA ratio, 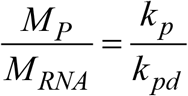. From Eq. 8 and Table 2, we find that 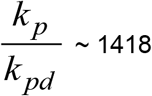 in both media, which agrees with previous estimations (~1832 in 27]).

Next, we used the fitted model to predict (using Eqs. 8 to 11) the influence of promoter occupancy (*ω*) on the *M_P_*, 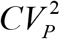 and *S_P_* of upstream and downstream promoters. We set d_TSS_ to 20 bp to represent promoters where ≤ 35, and to 100 bp to represent promoters with *d_TSS_* > 35. Then, for each cohort, we changed *ω* from 0.01 to 0.99 (i.e., nearly all possible values). In addition, we estimated these moments when *k_occlusion_*, *k_occupy_*, and *ω* are all set to zero (i.e., the two promoters do not interfere), for comparison.

From Fig. 7, a pair of tandem promoters can produce less proteins than a single promoter with the same parameter values, if *d_TSS_* ≤ 35, which makes occlusion possible. Meanwhile, if *d_TSS_* > 35, tandem promoters can only produce protein numbers in between the numbers produced by one isolated promoter and the numbers produced by two isolated promoters. In no case can two interfering tandem promoters produce more than two isolated promoters with equivalent parameter values. I.e., according to the model, the interference between tandem promoters cannot enhance production.

**Fig 7.**
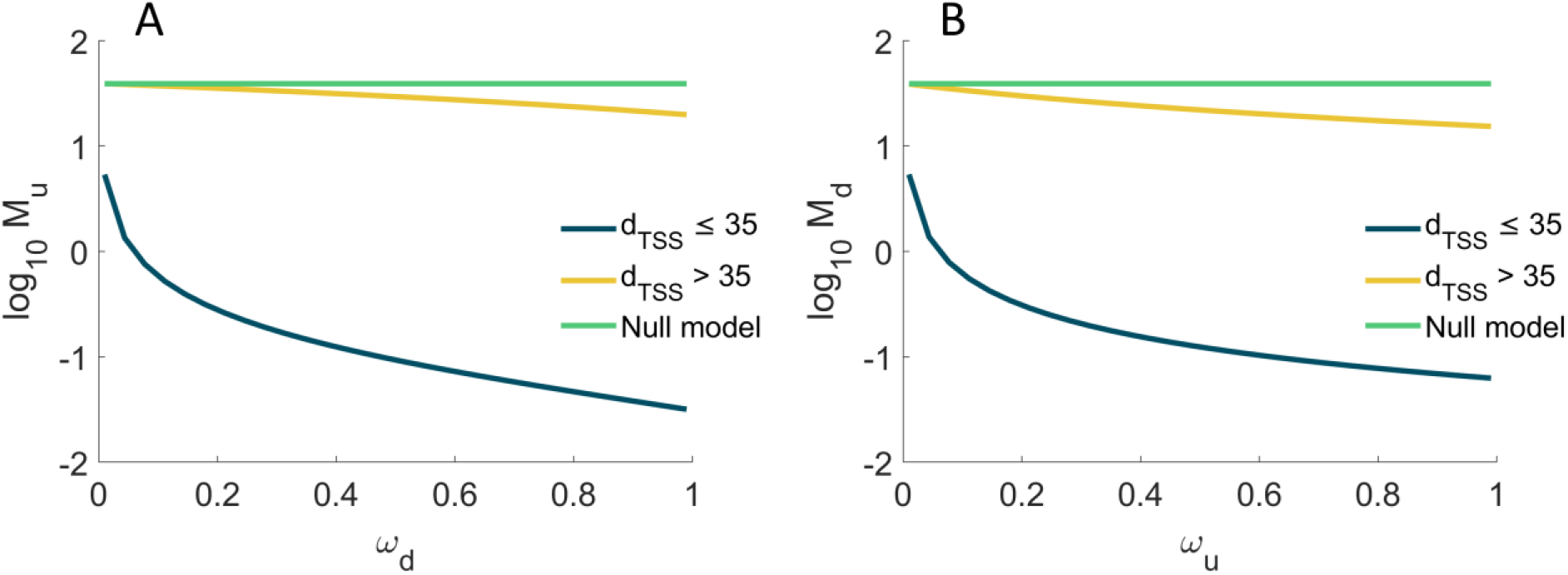
Mean protein numbers produced as a function of other promoter’s occupancy. *M_P_* of the single-cell distribution of the number of proteins produced (**A**) by the upstream promoter alone, and (**B**) by the downstream promoter alone. Results are shown as a function of the fraction of times that the upstream (0.01 ≤ *ω_u_* ≤ 0.99) and the downstream (0.01 ≤ *ω_d_* ≤ 0.99) promoter are occupied by RNAP. The null model is estimated by setting *k_occlusion_*, *k_occupy_*, and *ω* to zero.

Meanwhile, the kinetics of the upstream (Figs. 7A and S15A in the S2 Appendix) and downstream promoters (Figs. 7B and S15B in the S2 Appendix) only differ in that the downstream promoter is more responsive to *ω*.

Finally, consider that the model predicts that transcription interference should occur in tandem promoters, either due to occlusion if *d_TSS_* ≤ 35 occupancy or due to occupancy of the downstream promoter if *d_TSS_* > 35. Meanwhile, in single promoters, neither of these phenomena occurs. Thus, on average, two single promoters should produce more RNA and proteins than a pair of tandem promoters of similar strength. Using the genome wide data from [28] on protein expression levels during exponential growth we estimated the double of the mean expression level (it equals 183.8) of genes controlled by single promoters (section ‘Selection of natural genes controlled by single promoters’ in the S1 Appendix). Meanwhile, also using data from [28], the mean expression level of genes controlled by tandem promoters equals 148 (estimated from the 26 that they have reported on), in agreement with the hypothesis. Nevertheless, this data is subject to external variables (e.g., TF interference). A definitive test would require the use of synthetic constructs, lesser affected by external influences.

### Regulatory parameters of promoter occupancy and occlusion

Since the occupancy, ω, of each of the tandem promoters is responsible for transcriptional interference by occlusion and by RNAPs occupying the downstream promoter, we next explored the biophysical limits of *ω*. Eqs. 6a and 6b define the occupancies of the upstream and downstream promoters, *ω_u_* and *ω_d_*, respectively. For simplicity, here we refer to both of them as *ω*. Fig. 8A shows that *ω* increases with the rate of RNAP binding (*k_bind_* · [*R*]), but only within a certain range of (high) values of the time from binding to elongating 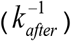. I.e., RNAPs need to spend a significant time in OC, if they are to cause interference, which is expected. Similarly, *ω* changes with 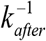, but only for high values of *k_bind_* · [*R*]. I.e., if it’s rare for RNAPs to bind, the occupancy will necessarily be weak.

**Fig 8.**
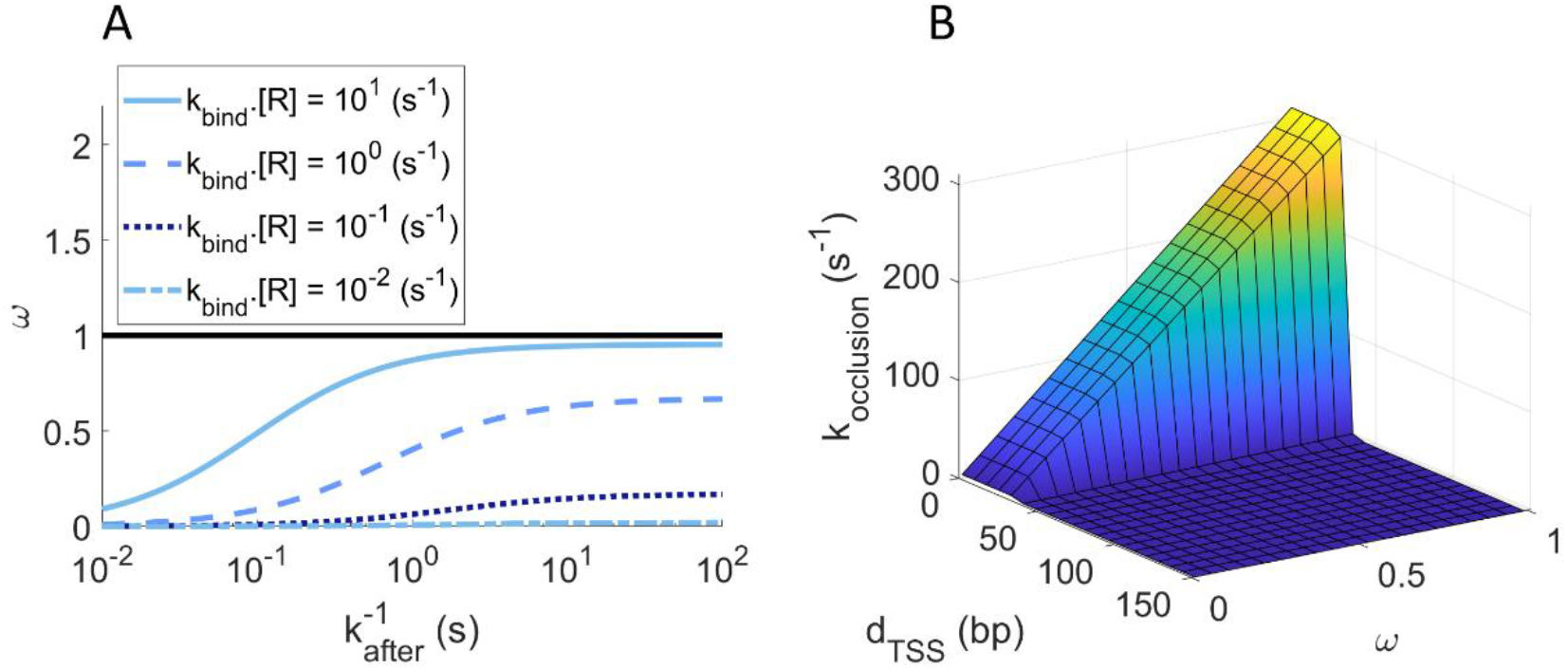
Promoter occupancy *ω* estimated for the step model. (A) *ω* as a function of the rate constant for a *free* RNAP to bind to the *unoccupied* promoter (*k_bind_* · [*R*]) and of the time for that RNAP to start elongation after commitment to OC, 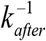. The horizontal black line at *ω* = 1, is the maximum fraction of time that the promoter can be occupied (i.e., the maximum promoter occupancy). (B) *k_occlusion_* plotted as a function of *ω* and *d_TSS_*. Since *k_occlusion_* increases with *ω* if and only if *d_TSS_* ≤ 35, it renders the simultaneous occupation of both TSS’s impossible.

In detail, from Fig. 8A, *ω* can change significantly within 10^−2^ < *k_bind_* × *[R]* < 10 s^−1^ and 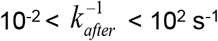. For these ranges, we expect RNA production rates (*k_r_*, equations 5a, 5b, 6b, 7 and 9) to vary from ~10^−5^ (if *d_TSS_* ≤ 35) and ~10^−4^ (if *d_TSS_* > 35) until 10 s^−1^. In agreement, in *E. coli*, promoters have RNA production rates from ~10^−3^ to 10^−1^ s^−1^ when induced [20–21, 39, 51–52]and ~10^−4^ to 10^−6^ s^−1^ when non-fully active [28]. Thus, *ω* can differ within realistic intervals of parameter values.

Next, we estimated *k_occlusion_*, the rate at which a promoter occludes the other as a function of *d_TSS_* and *ω* using Equations 6a and 6b. *k^max^* is shown in Table 3. To model *I(d_TSS_)* we used the step function in Table 1. Overall, *k_occlusion_* changes linearly with *ω*, when and only when *d_TSS_* ≤ 35 (Fig. 8B).

### State space of the single cell statistics of protein numbers of tandem promoters

We next studied how much the single-cell statistics of protein numbers (*M_P_*, 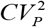, and *S_P_*) of the upstream, ‘u’, and downstream, ‘d’, promoters changes with *ω_u_*, *ω_d_* and *d_TSS_*. Here, *ω_u_* and *ω_d_* are increased from 0 to 1 by increasing the respective *k_bind_* (Eqs. 6a and 6b).

From Fig. 9A, if *d_TSS_* ≤ 35 bp, reducing *ω_d_* while also increasing *ω_u_* is the most effective way to increase *Mu*, since this increases the number of RNAPs transcribing from the upstream promoter that are not hindered by RNAPs occupying the downstream promoter. If *d_TSS_* > 35 bp, the occupancy the downstream promoter, *ω_d_*, becomes ineffective.

**Fig 9.**
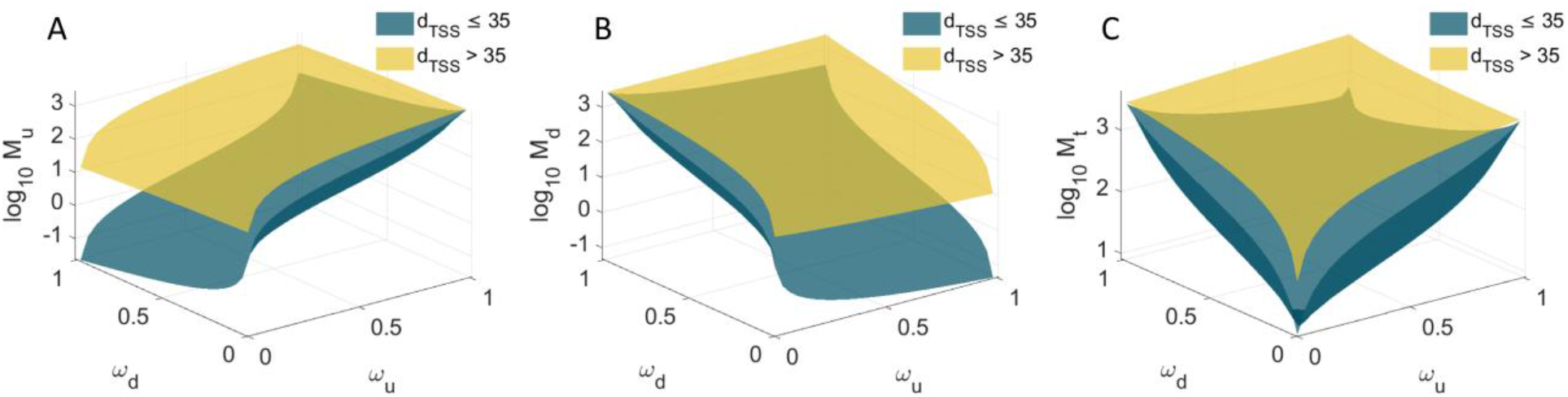
Mean protein expression as a function of both promoters’ occupancy. Expected mean protein numbers due to the activity of: (**A**) the upstream promoter alone, (**B**) the downstream promoter alone, and (**C**) both promoters. M_P_ is shown as a function of the fraction of times that the upstream (0 ≤ *ω_u_* ≤ 1) and the downstream (0 ≤ *ω_d_* ≤ 1) promoters are occupied by RNAP, when *d_TSS_* > 35 (yellow) and *d_TSS_* ≤ 35 (dark green) bp.

Oppositely, from Fig. 9B, if *d_TSS_* ≤ 35 bp, increasing *ω_d_* while also decreasing *ω_u_*, is the most effective way to increase M_d_ since this increases the number of RNAPs transcribing from the downstream promoter does not interfere by RNAPs elongating from the upstream promoter. If *d_TSS_* > 35 bp, the occupancy the upstream promoter, *ω_u_*, becomes ineffective.

Finally, from Fig. 9C, regardless of *d_TSS_*, for small *ω_d_* and *ω_d_*, as the occupancies increase, *M_t_* increases quickly and in a non-linear fashion. However, as both *ω_d_* and *ω_u_* reach high values, *M_t_* decreases for further increases, if *d_TSS_* ≤ 35 bp. Instead, if *d_TSS_* > 35 bp, Mt appears to saturate.

From Figs. S16 in the S2 Appendix, 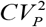 and *S_P_* behave inversely to *M_P_*.

Relevantly, in all cases, the range of predicted protein numbers (Fig. 9C1) are in line with the empirical values (~10^−1^ to 10^3^ proteins per cell) (Fig. 4D).

## Discussion

*E. coli* genes controlled by tandem promoters have a relatively high mean conservation level (0.2, while the average gene has 0.15, with a p-value of 0.009), suggesting that they play particularly relevant biological roles (section ‘Gene Conservation’ in the S1 Appendix). From empirical data on single-cell protein numbers of 30 *E. coli* genes controlled by tandem promoters, we found evidence that their dynamics is subject to RNAP interference between the two promoters. This interference reduces the mean single-cell protein numbers, while increasing its CV^2^ and skewness, and can be tuned by *ω*, the promoters’ occupancy by RNAP, and by *d_TSS_*. Since both of these parameters are sequence dependent [21, 31] the interference should be evolvable. Further, since *ω* of at least some of these genes should be under the influence of their several input TFs, the interference has the potential to be adaptive.

We proposed models of the dynamics of these genes as a function of *ω* and *d_TSS_*, using empirically validated parameter values. In our best fitting model, transcription interference is modelled by a step function of *d_TSS_* (instead of gradually changing with *d_TSS_*), since the only detectable differences in dynamics with changing *d_TSS_* were between tandem promoters with *d_TSS_* ≤ 35 and *d_TSS_* > 35 nucleotides (the latter cohort of genes having higher mean expression and lower variability). We expect that causes this difference tangible is the existence of the OC formation. In detail, the OC is a long-lasting DNA-RNAP formation that occupies that strict region of DNA at the promoter region [24, 31]. As such, occlusion should share these physical features. Because of that, when *d_TSS_* ≤ 35, an RNAP bound to TSS always occludes the other TSS, significantly reducing RNA production. Meanwhile, if *d_TSS_* > 35, interference occurs when an RNAP elongating from the upstream promoter is obstructed by an RNAP occupying the downstream promoter.

Meanwhile, contrary to *d_TSS_*, if one considers realistic ranges of the other model parameters, it is possible to predict a very broad range of accessible dynamics for tandem promoter arrangements. This could explain the observed diversity of single-cell protein numbers as a function of *d_TSS_* (Fig 6). At the evolutionary level, such potentially high range of dynamics may provide high evolutionary adaptability and thus, it may be one reason why genes controlled by these promoters are relatively more conserved.

One potentially confounding effect which was not accounted for in this model is the accumulation of supercoiling. Closely spaced promoters may be more sensitive to supercoiling buildup than single promoters [53–55]. If so, it will be useful to extend the model to include these effects [26]. Using such model and measurements of expression by tandem promoters when subject to, e.g. Novobiocin [56], may be of use to infer kinetic parameters of promoter locking due to positive supercoiling build-up.

Other potential improvements could be expanding the model to tandem arrangements other than I and II (Fig 1), to include a third form of interference (transcription elongation of a nearby gene).

One open question is whether placing promoters in tandem formation increases the robustness of downstream gene expression to perturbations (e.g., fluctuations in the concentrations of RNAP or TF regulators). A tandem arrangement likely increases the robustness to perturbations which only influence one of the promoters. Another open question is why several of the 102 tandem promoters with arrangements I and II appeared to behave independently from their input TFs (according to the RNA-seq data), albeit having more input TFs (1.62 on average) than expected by chance (the average *E. coli* gene only has 0.95). As noted above, we hypothesize that these input TFs may become influential in conditions other than the ones studied here.

Here, we also did not consider any influence from the phenomenon of “RNAP cooperation” [57]. This is based on this being an occurrence in elongation, and we expect interactions between two *elongating* RNAPs to rarely affect the interference between tandem promoters [9]. However, potentially, it could be of relevance in the strongest tandem promoters.

Finally, a valuable future study on tandem promoters will require the use of synthetic tandem promoters (integrated in a specific chromosome location) that systematically differ in promoter strengths and nucleotide distances. This would allow extracting parameter values associated to promoter interference to create a more precise model than the one based on the natural promoters (which is influenced by TFs, etc). Similarly, measuring the strength of individual natural promoters would contribute to this effort.

Overall, our model, based on a significant number of natural tandem promoters whose genes have a wide range of expression levels, should be applicable to the natural tandem promoters not observed here (at least of arrangements I and II), including of other bacteria, and to be accurate in predicting the dynamics of synthetic promoters in these arrangements.

Currently, predicting how gene expression kinetics change with the promoter sequence remains challenging. Even single- or double-point mutants of known promoters behave unpredictably, likely because the individual sequence elements influence the OC and CC in a combinatorial fashion. Consequently, the present design of synthetic circuits is usually limited to the use of a few promoters whose dynamics have been extensively characterized (Lac, Tet, etc.). This severely limits present synthetic engineering.

We suggest that a promising methodology to create new synthetic genes with a wide range of predictable dynamics is to assemble well-characterized promoters in a tandem formation, and to tune their target dynamics using our model. Specifically, for a given dynamics, it is possible to invert the model and find a suitable pair of promoters with known occupancies and corresponding *d_TSS_* (smaller or larger than 35), which achieve these dynamics. A similar strategy was recently proposed in order to achieve strong expression levels [58]. Our results agree and further expand on this by showing that the mean expression level can also be reduced and expression variability can further be fine-tuned.

Importantly, this can already be executed, e.g., using a library of individual genes whose expression can be measured [28]. From this library, we can select any two promoters of interest and arrange them as presented here, in order to obtain a kinetics of expression as close as possible to a given target. Note that these dynamics have a wide range, from weaker to stronger than that of either promoter (albeit no stronger than their sum, Fig 9C1-C3). Given the number of natural genes whose expression is already known and given the present accuracy in assembling specific nucleotide sequences, we expect this method to allow the rapid engineering of genes with desired dynamics with an enormous range of possible behaviours. As such, these constructs could represent a recipe book for the components of gene circuits with predictable complex kinetics.

## Materials and Methods

Using information from RegulonDB v10.5 as of 30^th^ of January 2020, we started by searching natural genes controlled by two promoters (Section ‘Selection of natural genes controlled by tandem promoters’ in the S1 Appendix). Next, we studied their evolutionary conservation and ontology (Sections ‘Gene conservation’ and ‘Gene Ontology’ in the S1 Appendix) and analysed their local topological features within the TFN of *E. coli* (Section ‘Network topological properties’ in the S1 Appendix).

RNA-seq measurements were conducted in two points in time (Section ‘RNA-seq measurements and data analysis’ in the S1 Appendix), to obtain fold changes in RNA numbers of genes controlled by tandem promoters with arrangements I and II, their input TFs, and their output genes (Fig 1). We used this data to search for relationships between input and output genes.

Next, a model of gene expression was proposed, and reduced to obtain an analytical solution of the single-cell protein expression statistics of tandem promoters (Sections ‘Derivation of mean protein expression of the model’ and ‘Derivation of *CV^2^* and skewness of protein expression of the model’ in the S1 Appendix). This analytical solution was compared to stochastic simulations conducted using the simulator SGNS2. (Section ‘Stochastic simulations for the step inference model’ in the S1 Appendix).

We collected single-cell flow-cytometry measurements of 30 natural genes controlled by tandem promoters (Section ‘Flow-cytometry and data analysis’ in the S1 Appendix) to validate the model. For this, first, from the original data, we subtracted the cellular background fluorescence (Section ‘Subtraction of background fluorescence from the total protein fluorescence’ in the S1 Appendix). Then, we converted the fluorescence intensity into protein numbers (Section ‘Conversion of protein fluorescence to protein numbers in the S1 Appendix). From this we obtained empirical data on *M, CV^2^, and S* of the single-cell distributions of protein numbers in two media (Sections ‘Media and chemicals’ and ‘Strains and growth conditions’ in the S1 Appendix). Flow-cytometry measurements were also compared to microscopy data, supported by image analysis (Section ‘Microscopy and Image analysis’ in the S1 Appendix), for validation.

Comparing the data from RegulonDB (30.01.2020) used here, with the most recent (21.07.2021), we found that the numbers of genes controlled by tandem promoters of arrangements I and II differed by ~4% (from 102 to 98). Regarding those whose activity was measured by flow-cytometry, this difference is ~3% (30 to 31). Globally, 163 TF-gene interactions differed (~3.4%) while for the 98 genes controlled by tandem promoters of arrangements I and II, only 10 TF-gene interactions differ (~2.7%). Finally, globally the numbers of TUs differed by ~1%, promoters by ~0.6%, genes by ~1%, and terminators by ~15% (which did not affect the genes studied, as they changed by ~4% only). These small differences should not affect our conclusions.

Finally, a data package [59] is provided in Dryad with flow-cytometry and microscopy data and codes used.

## Supporting information

Supporting Results

Gene Ontology table

Tandem promoter genes' protein statistics table

Single promoter genes' protein statistics table

## Supporting Information

**S1 Appendix. Extended Methods and Materials.** (PDF)

**S2 Appendix. Supporting Figures**. (PDF)

**S3 Appendix. Supporting Tables**. (PDF)

**S4 Appendix. Supporting Results**. (PDF)

**S5 Table. Gene Ontology**. Overrepresentation tests using the PANTHER Classification System. List of biological processes which are overrepresented using Fisher’s exact tests are shown. (Excel)

**S6 Table. Protein statistics**. Statistics of single-cell distributions of protein fluorescence of genes controlled by tandem promoters as measured by flow-cytometry in 1X and 0.5X diluted M9 media conditions. (Excel)

**S7 Table. Protein statistics**. Statistics of single-cell distributions of protein fluorescence of genes controlled by single promoter as measured by flow-cytometry in 1X M9 media condition. (Excel)

## Acknowledgements

The authors thank Jason Lloyd-Price for proof-reading and editing the text. The authors also thank all three referees for their valuable suggestions.

