## Extended materials and methods for "Analytical kinetic model of native tandem promoters in *E. coli*"

### S1 Appendix: Extended Materials and Methods

#### Selection of natural genes controlled by tandem promoters

We define a pair of tandem promoters as two promoters in a head-to-tail formation transcribing the same gene, as in [1]. In order to find them in the genome of *E. coli*, from RegulonDB, we obtained the lists of all known transcription units (TUs), promoters (defined as stretches of 60 upstream and 20 downstream nucleotide sequences from a TSS), gene sequences, TFs, and terminators [2].

From the list of TUs (3560), we extracted all genes (510) under the control of two and only two promoters in tandem formation with known TSS and DNA strand (information from the promoters' list). Then, we calculated the nucleotide distance between their pair of TSSs ($d_{TSS}$) and obtained the start and end positions of their sequence in the DNA. As a side note, we found additional 321 genes controlled by more than two promoters in tandem formation, which are not accounted for as they are not included in the model, for simplicity.

Next, we removed all genes with another gene or promoter sequence (associated to a TU) located in the opposing strand anywhere between the start of the upstream promoter and the end of the gene sequence (186 out of 510) since their dynamics may be subject to interference from convergent RNAPs [1,3,4]

Out of the remaining 324 genes, only 152 are in the first position of a TU or in a TU with only one gene. Since evidence suggests that the existence of multiple genes in a TU influences their transcription significantly, due to premature terminations, distance to the promoter etc. [5,6], we opted for keeping only those 152 genes. Subsequently, from the list of terminators, we obtained their start and end positions and DNA strand and filtered out (9 out of 152) genes with a terminator sequence in between the beginning of the upstream promoter and the end of the gene sequence, due to potential enhanced premature terminations. Finally, from these, we only considered promoter pairs (102 out of the 143 genes) such that no gene is coded in the regions containing them or the space in between them (Fig 1), so that elongation of other genes do not perturb their transcription.

Finally, of these 102 genes, we measured the expression levels at the single-cell level of 30 of them (Table S1 in S3 Appendix) using a YFP strain library [7]. These genes are of the categories 'I' (9 genes) and 'II' (21 genes) in Figure1. Their $d_{TSS}$'s range from 84 to 173, and from 3 to 73 nucleotides, respectively.

#### Selection of natural genes controlled by single promoters

To select natural genes controlled by single promoters in the genome of *E. coli,* from RegulonDB, we obtained the lists of all known transcription units (TUs), promoters, gene sequences and terminators [2]. From the list of TUs (3560), we extracted all genes (1760) under the control of one and only one promoter with known TSS and DNA strand (information from the promoters' list). Next, we filtered out all genes with another gene or promoter sequence (associated to a TU) located in the opposing strand anywhere between the start of the promoter and the end of the gene sequence (446 out of 1760) since their dynamics may be subject to interference from convergent RNAPs [1,3,4] Out of the remaining 1314 genes, only 649 are in the first position of a TU or in a TU with only one gene and no other promoter sequence (associated to another TU) between the promoter and the end of the gene of interest. Since evidence suggests that the existence of multiple genes in a TU influences their transcription significantly, due to premature terminations, distance to the promoter etc. [5,6], we opted for keeping only those 649 genes. Subsequently, from the list of terminators, we obtained their start and end positions and DNA strand and filtered out (36 out of 649) genes with a terminator sequence in between the promoter and the end of the gene sequence, due to potential enhanced premature terminations. Finally, of these 613 genes, we obtained data on the expression levels of 126 genes from [7], which we used to compare expression levels of genes controlled by tandem promoters and genes controlled by single promoters.

Meanwhile, for purposes of validating the scaling factor between protein fluorescence and numbers, of these 613 genes, we measured the expression levels at the single-cell level of 10 of them, randomly selected (Table S2 in S3 Appendix) [7].

#### Gene Conservation

From a list of 5443 reference bacterial genomes [8], we used the Rentrez package [9] to obtain which genes are present in each genome. Next, we removed those genomes without gene entries (1310). Using the remaining genomes, we estimated the evolutionary conservation of each gene in the genome of MG1655 (GCF_000005845.2_ASM584v2), including those controlled by tandem promoters, by the ratio between the number of genomes where the gene is present and the total number of genomes considered. Supplementary Fig. S17 in S2 Appendix shows the conservation levels as a function of $d_{TSS}$ of the tandem promoters controlling the genes' expression.

#### Gene Ontology (GO)

For gene ontology representations, we performed overrepresentation tests using the PANTHER Classification System [10], which finds statically significant overrepresentations using Fisher's exact tests. For p-values < α (here set to 0.05), the null hypothesis that there are no associations between the gene cohort and the corresponding GO of the biological process is rejected, which we interpret as the gene cohort being associated with corresponding GO of the biological process.

#### Network topological properties

By 'network topological property' we refer to some feature of a gene that is related to how that gene is integrated with the network formed by TFs linking genes. We used *Cytoscape* [11] to extract these features for the genes controlled by tandem promoters from the *known* transcription factor (TF) network of *E. coli*, using information from RegulonDB v10.5 on all known transcription factors (TFs) and their binding sites [2].

Next, for the two cohorts of genes with $d_{TSS}$ larger or not than 35 bps, based on definitions in [12], we calculated (Table S3 in S3 Appendix) the mean and standard error of each cohort's average shortest path length (minimum number of edges between pairs of genes), clustering coefficient (fraction of input nodes to a node that are also linked), eccentricity (maximum non-infinite shortest path length between the node and another node in the network), edge count (number of edges/nodes that are connected to the node), indegree (number of incoming edges), neighbourhood connectivity (average connectivity of all nearest neighbours), and outdegree (number of outgoing edges).

For each feature, we also obtained a p-value, which is the probability that the genes of the cohort have a smaller mean than the mean from all genes of *E. coli*. This probability is estimated from $10^5$ cohorts assembled from random samples from all genes with replacement, using a non-parametric bootstrap method. The sample size is equal to the size of the cohort being compared with.

### Media and chemicals

Measurements were performed in Luria-Bertani (LB) and M9 media (standard and diluted). The chemicals, such as tryptone, sodium chloride, agarose, MEM amino acids (50X), MEM Vitamin solution (100X), Glucose and antibiotic chloramphenicol, etc. were purchased from Sigma Aldrich. Yeast extract was purchased from Lab M (Topley House, Bury, Lancashire, UK). The components of LB medium were 10 g tryptone, 10 g NaCl, and 5 g yeast extract in 1000 mL distilled water. For M9 medium, the components were 1x M9 Salts, 2 mM MgSO4, 0.1 mM CaCl2; 5x M9 Salts with 34 g/L Na2HPO4, 15 g/L KH2PO4, 2.5 g/L NaCl, 5 g/L NH4Cl supplemented with 100X vitamins, 0.2% Casamino acids and 0.4% glucose. We also used '0.5X' and '0.25X' media by diluting the M9 medium to 1:1 and to 1:3 respectively, using autoclaved distilled water [13-16].

### Strains and growth conditions

To measure RNA polymerase (RNAP) levels at different medium, we used the RL1314 strain with RpoC endogenously tagged with GFP (generously provided by Robert Landick), which was engineered from the W3110 strain (used here to measure background fluorescence).

To measure single-cell protein levels of genes controlled by tandem promoters, we used genes endogenously tagged with the YFP coding sequence from the YFP fusion library [7]. These were purchased from the *E. coli* genetic stock center (CGSC) of Yale University, U.S.A. (Table S2 in S3 Appendix), which has wild type MG1655 cells as the reference genome (and thus was used to measure cellular background fluorescence). Measurements of protein levels using this library are expected to be precise for a wide range of expression levels, given evidence for strong correlation in single gene expression levels when measured by RNA-fish, RNA-seq, mass spectrometry and flow cytometry (taken using the YFP library) [7]. The lesser accurate estimations occur for the weakest expressing genes

[7][17], due to their values being near the level of cellular autofluorescence. For this reason as well, we do not consider all of the 30 genes in our analysis as described in the Results section.

From a glycerol stock (-80°C), cells were streaked on LB agar plates with the appropriate antibiotics and incubated at 37°C overnight. From the plates, a single colony was picked, inoculated in LB medium and supplemented with appropriate antibiotics and incubated at 30°C overnight with shaking at 250 rpm. Next, overnight cultures were diluted into freshly prepared tailored media (see 'Media and Chemicals'), with appropriate antibiotics with an $O.D_{600}$ of 0.03 (Optical Density, 600 nm; Ultrospec 10, Amersham biosciences, UK) and allowed to grow at 30°C with shaking at 250 rpm until reaching the mid-exponential phase ($O.D_{600}$ ~0.4-0.5). At this stage, measurements of protein levels were conducted using flow-cytometry and/or microscopy.

### Growth curves

Growth curves were measured by $O.D_{600}$ using a spectrophotometer (Ultrospec 10; GE Healthcare). From the overnight culture, cells were diluted (1:10000) into the respective fresh media and allowed to grow while shaking (250 rpm). O.D.'s were recorded for 450 min. every 30 min. We performed 3 biological replicates for each condition. We found negligible variability between replicates. The results shown are the averages and standard error of the mean.

### Microscopy and image analysis

When reaching the mid-exponential growth phase, cells were pelleted by centrifugation (10000 rpm for 1 min), and the supernatant was discarded. The pellet was re-suspended in 100 µL of the remaining medium Next, 3 µL of cells were placed in between 2% agarose gel pad and a coverslip and imaged using a confocal microscopy with a 100X objective. The fluorescence was measured with a 488 nm laser and a 514/30 nm emission filter. Phase-contrast images were simultaneously acquired for purposes of segmentation and to assess health, morphology, and physiology.

Using the software *CellAging* [18], from phase contrast images, we segmented cells semi-automatically, correcting errors manually. Next, phase-contrast and corresponding fluorescence images were aligned to extract single-cell fluorescence intensities (example image in Fig 4B). We then performed background subtraction, i.e., from each cell's total fluorescence we subtracted the mean fluorescence of control cells, not expressing YFP.

### RNA-seq measurements and data analysis

We searched for correlations between the LFCs over time of genes controlled by tandem promoters ('Tg') and the LFCs over time of their output genes ('Og') as well as their input genes ('Ig').

Given known rates of RNA and protein production and degradation in *E. coli* [7, 19-22], we expect changes in RNA numbers to take at least 60 min. on average, to propagate to protein numbers. Thus, we performed RNA-seq of cells in exponential growth phase at moments '0 min', and then 20 and 180 mins. later. We then calculated LFCs between 0 and 20 min, and between 0 and 180 min.

Specifically, to assess if LFCs in Ig propagate to Tg, we compared changes in Ig between moments 0 and 20 min, with changes in Tg between moments 0 and 180 min. Similarly, to assess LFCs in Tg propagate to Og, we compared changes in Tg between moments 0 and 20 min, with changes in Og between moments 0 and 180 min. Results are shown in Figs S4A and S4B in S2 Appendix.

#### Sample preparation

For RNA-seq experiments, single colonies of K12 MG1655 cells were picked from LB Agar plates and inoculated into 5 ml of LB medium. Cultures were grown overnight with shaking at 250 rpm. Next, these cultures were diluted to $O.D_{600}$ of 0.05 in fresh LB medium and incubated, with a 250 rpm agitation. RNA-seq was performed over time (0, 20 and 180 min). Total RNA from 3 independent biological replicates in each medium was extracted using RNeasy kit (Qiagen). RNA was treated twice with DNase (Turbo DNA-free kit, Ambion) and quantified using Qubit 2.0 Fluorometer RNA assay (Invitrogen, Carlsbad, CA, USA). Total RNA amounts were determined by gel electrophoresis, using a 1% agarose gel stained with SYBR safe (Invitrogen). RNA was detected using UV with a Chemidoc XRS imager (Biorad).

Sequencing was performed by GENEWIZ, Inc. (Leipzig, Germany). The RNA integrity number (RIN) was obtained with the Agilent 4200 TapeStation (Agilent Technologies, Palo Alto, CA, USA). Ribosomal RNA depletion was performed using Ribo-Zero Gold Kit (Bacterial probe) (Illumina, San Diego, CA, USA). RNA-seq libraries were constructed using NEBNext Ultra RNA Library Prep Kit (NEB, Ipswich, MA, USA). Sequencing libraries were multiplexed and clustered on 1 lane of a flow-cell. Samples were sequenced using a single-index, 2x150 bp paired-end (PE) configuration on an Illumina HiSeq instrument. Image analysis and base calling were conducted with HiSeq Control Software (HCS). Raw sequence data (.bcl files) were converted into fastq files and de-multiplexed using Illumina bcl2fastq v.2.20. One mismatch was allowed for index sequence identification.

#### Data analysis

RNA-seq data analysis pipeline was: i) RNA sequencing reads were trimmed with Trimmomatic [23] v.0.39) to remove possible adapter sequences and nucleotides with poor quality. ii) Trimmed reads were mapped to the reference genome, *E. coli* MG1655 (NC_000913.3), using the using the STAR aligner v.2.5.2b, which outputs BAM files [24]. iii) Then, '*featureCounts'* from the Rsubread R package (v.1.34.7) was used to calculate unique gene hit counts [25]. iv) These counts were used for the differential expression analysis. Genes with less than 5 counts in more than 3 samples, and genes whose mean counts are less than 10 were removed from further analysis. We used the DESeq2 R package (v.1.24.0) [26] to compare gene expression between groups of samples and calculate p-values and $\log_2$ of fold changes using Wald tests (function *nbinomWaldTest*). P-values were adjusted for multiple hypotheses testing (Benjamini–Hochberg, BH procedure, [27]).

#### Flow-cytometry and data analysis

We measured single-cell fluorescence using a ACEA NovoCyte Flow Cytometer (ACEA Biosciences Inc., San Diego, USA). Upon reaching the mid-exponential phase (OD~0.4-0.5), cells were diluted (1:10000) into 1 mL of phosphate buffer saline (PBS) solution and vortexed for 5 s. For a single run, 50000 events were collected at a flow rate of 14 µL/minute and a core diameter of 7.7 mm using the Novo Express software using a blue laser (488 nm) for excitation. We obtained the height of the fluorescein isothiocyanate channel (FITC-H) (530/30 nm filter). A PMT voltage of 600 volts was set for FITC. To avoid background signal from particles smaller than bacteria, the detection threshold was set to 5000 for FSC-H analyses. Three biological replicates were performed per condition.

We applied unsupervised gating [28] to the flow-cytometry data, setting the fraction of single-cell events used in the analysis, α, to 0.99. We proved to be enough to remove non-cell events due to debris, doublets, fragments, cell clumps, and other undesired events. Reducing α did not change the results qualitatively.

To remove outliers from the flow-cytometry distributions, we applied secondary gating. In detail, we sorted the data based on FITC-H values and calculated the difference between consecutive samples. Then, we obtained the indices of those differing by more than 10000 (approximately 10 times the mean fluorescent level observed). Next, we obtained the minimum of those indices to define the upper bound. Finally, values above this index were considered an outlier and discarded. In all measurements, never more than 10000 events were discarded, thus, more than 40000 were used for the analysis.

#### Subtraction of background fluorescence from total protein fluorescence in flow-cytometry

First, we collected mean background fluorescence from distributions of cells not carrying YFP. Then we measured the distributions of fluorescence of cells carrying the protein tagged with YFP. Having this, the protein fluorescence 'g' of a gene is obtained by subtracting mean background fluorescence 'bg' from the (total 'T') measured fluorescence. For the mean (M) protein fluorescence from a cell population, we write:

$$M\left(g\right)=M(T)-M(bg) \tag{S1}$$

Similarly, the variance 'Var' is obtained by:

$$Var(g)=Var(T)-Var(bg) \tag{S2}$$

The $CV^2$ of the distribution protein fluorescence of a gene after background subtraction is:

$$CV^2(g) = \frac{Var(g)}{M(g)^2} \quad \text{(S3)}$$

Finally, the third moment of protein fluorescence and the skewness after background subtraction are given by:

$$\mu_3(g) = \mu_3(T) - \mu_3(bg) \quad \text{(S4)}$$

$$S(g) = \frac{\mu_3(g)}{Var(g)^{\frac{3}{2}}} \quad \text{(S5)}$$

After background subtraction, any genes with negative means, variance or third moment, will not be included in the data (except in Fig S6 in S2 Appendix for illustrative purposes).

### Conversion of protein fluorescence into protein numbers

To convert protein fluorescence into protein numbers, we made a correlation plot between the mean protein fluorescence measured in our lab (after background subtraction) and the mean protein numbers reported in [7] for the same genes. We fitted a line to the data points by forcing the intercept with the Y axis to be at zero. The slope of the fitted line is used as a scaling factor (~0.09) with an $R^2$ value of 0.68 (Fig 4D). For protein fluorescence to protein numbers correction only the mean gets changed whereas the normalised moments $CV^2$ and S remain unchanged.

### Analytical model of mean RNA levels controlled by a single promoter in the absence of a closely spaced promoter

From Reactions 1c1 and 1a4 in the main manuscript, for an isolated promoter, one would have:

$$P_{free} \underset{k_{unbind}}{\overset{k_{bind}\cdot[R]}{\rightleftarrows}} P_{occupied} \xrightarrow{k_{after}} P_{free} + R_{elong} \quad \text{(S6)}$$

$$R_{elong} \xrightarrow{k_{elong}} RNA \quad \text{(S7)}$$

At steady state $P_{occupied}$ is:

$$\frac{dP_{occupied}}{dt} = P_{free} \times k_{bind} \cdot [R] - P_{occupied} \times \left(k_{unbind} + k_{after}\right) = 0 \quad \text{(S8)}$$

$$P_{free} = P_{occupied} \cdot \frac{\left(k_{unbind} + k_{after}\right)}{k_{bind} \cdot [R]} \tag{S9}$$

Since necessarily:

$$P_{free} + P_{occupied} = 1 \tag{S10}$$

From equations S9 and S10:

$$P_{occupied} \cdot \left(1 + \frac{k_{unbind} + k_{after}}{k_{bind} \cdot [R]}\right) = 1 \tag{S11}$$

$$P_{occupied} = \frac{k_{bind} \cdot [R]}{k_{bind} \cdot [R] + k_{unbind} + k_{after}} \tag{S12}$$

Note that, by definition (main manuscript, equations 6a and 6b), the fraction of time that an RNAP is bound to the promoter, $\omega$, should equal $P_{occupied}$ in (S12). Meanwhile, at steady state, $R_{elong}$ becomes:

$$\frac{dR_{elong}}{dt} = P_{occupied} \times k_{after} - R_{elong} \times k_{elong} = 0 \tag{S13}$$

$$R_{elong} = \frac{P_{occupied} \times k_{after}}{k_{elong}} \tag{S14}$$

From equations S12 and S14:

$$R_{elong} = \frac{k_{bind} \cdot [R]}{k_{bind} \cdot [R] + k_{unbind} + k_{after}} \times \frac{k_{after}}{k_{elong}} \tag{S15}$$

At steady state, the mean $RNA$ numbers, $M_{RNA}$, is:

$$\frac{dM_{RNA}}{dt} = R_{elong} \times k_{elong} - M_{RNA} \times k_{rd} = 0 \tag{S16}$$

From equations S15 and S16s:

$$M_{RNA} = \frac{k_{bind} \cdot [R]}{k_{bind} \cdot [R] + k_{unbind} + k_{after}} \times \frac{k_{after}}{k_{elong}} \times \frac{k_{elong}}{k_{rd}} \tag{S17}$$

$$M_{RNA} = \frac{k_{bind} \cdot [R]}{k_{bind} \cdot [R] + k_{unbind} + k_{after}} \times \frac{k_{after}}{k_{rd}} \tag{S18}$$

From S18, the RNA numbers at steady state do not depend on $k_{elong}$.

### Derivation of mean protein numbers at steady state produced by a pair of tandem promoters

For the upstream promoter, from (1c1), (1a3), and (1a4) in the main manuscript, at steady state:

$$\frac{d(RNA)}{dt} = R^{u}_{elong} \times k^{u}_{elong} \cdot (1 - \omega_d \cdot f) - RNA \times k_{rd} = 0 \tag{S19}$$

From this and equation 6b in the main manuscript:

$$RNA = \frac{k^{u}_{bind} \cdot [R]}{k^{u/d}_{occlusion} + k^{u}_{bind} \cdot [R] + k^{u}_{unbind} + k^{u}_{after}} \times \frac{k^{u}_{after} \cdot (1 - \omega_d \cdot f)}{k_{rd}} \tag{S20}$$

Meanwhile, for the downstream promoter, from reactions (2a1), (2a2), and (2a3) in the main manuscript, at steady state:

$$\frac{d(RNA)}{dt} = R^{d}_{elong} \times k^{d}_{elong} - RNA \times k_{rd} = 0 \tag{S21}$$

$$RNA = \frac{k^{d}_{bind} \cdot [R]}{k^{d/u}_{occlusion} + k^{d}_{occupy} + k^{d}_{bind} \cdot [R] + k^{d}_{unbind} + k^{d}_{after}} \times \frac{k^{d}_{after}}{k_{rd}} \tag{S22}$$

Having this, since at steady state the mRNA numbers produced by a pair of tandem promoters should equal the sum of RNA numbers from the upstream (S20) and downstream (S22) promoters, we have:

$$M_{RNA} = \left( \frac{k_{bind}^{u} \cdot [R] \times k_{after}^{u} \cdot (1 - \omega_d \cdot f)}{k_{occlusion}^{u/d} + k_{bind}^{u} \cdot [R] + k_{unbind}^{u} + k_{after}^{u}} + \frac{k_{bind}^{d} \cdot [R] \times k_{after}^{d}}{k_{occlusion}^{d/u} + k_{occupy} + k_{bind}^{d} \cdot [R] + k_{unbind}^{d} + k_{after}^{d}} \right) \cdot \frac{1}{k_{rd}} \quad \text{(S23)}$$

Thus, the mean protein numbers is:

$$M_P = M_{RNA} \cdot \frac{k_p}{k_{pd}} \quad \text{(S24)}$$

If the upstream and downstream promoters have similar strengths, i.e., if $k_{bind}^{d} \approx k_{bind}^{u}$, $k_{unbind}^{d} \approx k_{unbind}^{u}$, and $k_{after}^{d} \approx k_{after}^{u}$, we can expect that: $\omega_d = \omega_u$, $k_{occlusion}^{d/u} = k_{occlusion}^{u/d}$. If so, the equation above becomes:

$$M_P = \left( \frac{k_{bind} \cdot [R] \times k_{after} \cdot (1 - \omega_d \cdot f)}{k_{occlusion} + k_{bind} \cdot [R] + k_{unbind} + k_{after}} + \frac{k_{bind} \cdot [R] \times k_{after}}{k_{occlusion} + k_{occupy} + k_{bind} \cdot [R] + k_{unbind} + k_{after}} \right) \cdot \frac{k_p}{k_{rd} \cdot k_{pd}} \quad \text{(S25)}$$

Here, the symbols "u" and "d" are removed, as they no longer imply potentially different amounts. Having this, let $k_r$ be the effective transcription rate constant of a pair of tandem proteins. It should equal:

$$k_r = \left( \frac{k_{bind} \cdot [R] \times k_{after} \cdot (1 - \omega_d \cdot f)}{k_{occlusion} + k_{bind} \cdot [R] + k_{unbind} + k_{after}} + \frac{k_{bind} \cdot [R] \times k_{after}}{k_{occlusion} + k_{occupy} + k_{bind} \cdot [R] + k_{unbind} + k_{after}} \right) \quad \text{(S26)}$$

Thus, from equation S25 and S26:

$$M_P = \frac{k_r \cdot k_p}{k_{rd} \cdot k_{pd}} \quad \text{(S27)}$$

### CV2 and skewness of the distribution of single-cell protein numbers of model tandem promoters

The distributions of protein numbers in *E. coli* cells, can, in general, be well approximated by a Gamma or by a negative binomial distribution [7]. We assume here a negative binomial distribution. For a given number of events, if *r* is the number of failures, *p* is the probability of success per event, and an 'event' is an attempt to produce a protein, then the mean, variance, and skewness of the single-cell distribution of protein numbers should equal:

$$M_P = \frac{pr}{1-p} \tag{S28}$$

$$Var_P = \frac{pr}{(1-p)^2} \tag{S29}$$

$$S_P = \frac{1+p}{\sqrt{pr}} \tag{S30}$$

The relationship between the mean, $CV^2$ could be written as:

$$CV_P^2 = \frac{1}{M_P} \cdot \left( \frac{Var_P}{M_P} \right) \tag{S31}$$

Substituting (S28) and (S29) in (S31)

$$CV_P^2 = \frac{\left( \frac{1}{1-p} \right)}{M_P} \tag{S32}$$

Rewriting the above equation by assuming a scaling factor $C_1$ as:

$$C_1 = \frac{1}{1-p} \tag{S33}$$

$$CV_P^2 = \frac{C_1}{M_P} \tag{S34}$$

Taking $\log_{10}$ on both sides

$$\log_{10}\left(CV_P^2\right)=\log_{10}(C_1)-\log_{10}\left(M_P\right) \quad \text{(S35)}$$

From [17], $C_1$ is approximated as

$$C_1=\frac{M_P}{M_{RNA}}\cdot\frac{\frac{1}{\tau_p}}{\frac{1}{\tau_p}+\frac{1}{\tau_{RNA}}} \quad \text{(S36)}$$

$\tau_p=\frac{1}{k_{pd}}$ and $\tau_{RNA}=\frac{1}{k_{rd}}$ are the lifetimes of proteins and RNAs, respectively. The above equation is rewritten as:

$$C_1=\frac{k_p}{k_{pd}}\cdot\frac{k_{pd}}{k_{pd}+k_{rd}} \quad \text{(S37)}$$

$$C_1=\frac{k_p}{k_{pd}+k_{rd}} \quad \text{(S38)}$$

From (S28) and (S30), the relationship between the mean, skewness could be written as:

$$S_P=\frac{\frac{1+p}{\sqrt{1-p}}}{\sqrt{M_P}} \quad \text{(S39)}$$

The equation can be rewritten assuming constant $C_2$ as:

$$C_2=\frac{1+p}{\sqrt{1-p}} \quad \text{(S40)}$$

$$S_P=\frac{C_2}{\sqrt{M_P}} \quad \text{(S41)}$$

Taking $\log_{10}$ on both sides

$$\log_{10}\left(S_P\right)=\log_{10}\left(C_2\right)-\frac{1}{2}\cdot\log_{10}\left(M_P\right) \quad \text{(S42)}$$

The constants $C_1$ and $C_2$ are related as follows. From equation S33:

$$p = 1 - \frac{1}{C_1} \tag{S43}$$

Inserting S43 in S40:

$$C_2 = \frac{2 - \frac{1}{C_1}}{\sqrt{\frac{1}{C_1}}} \tag{S44}$$

The equation can be rewritten as

$$C_2 = 2\sqrt{C_1} - \frac{1}{\sqrt{C_1}} \tag{S45}$$

### Stochastic simulations for the step inference model

Stochastic simulations of the models were done using the stochastic gene network simulator SGNS2 [29]. These stochastic models were compared to the analytical solutions to assess how much variability can there be in $k_{bind} \cdot [R]$ without the analytical solution deviating too much.

First, to compare analytical and stochastic solutions, we set $d_{TSS}$ between 0 and 180 with an increment of 30. For each $d_{TSS}$, we calculated the occlusion rate constant ($k_{occlusion}$) for upstream and downstream promoters (Equations 5a and 5b in the main manuscript). The other parameters are listed in Tables 2 and 3 in the main manuscript. To obtain protein numbers at steady state, we have set the simulation time to $10^5$ seconds and performed 1000 runs per condition. From these runs, for each condition, we calculated the mean, $CV^2$ and skewness, along with their standard errors using bootstrapping ($10^4$ resampling with replacement). Additional runs would slightly decrease the deviation between the two solutions.
