## Supporting Figures for "Analytical kinetic model of native tandem promoters in *E. coli*"

### S2 Appendix: Supporting Figures

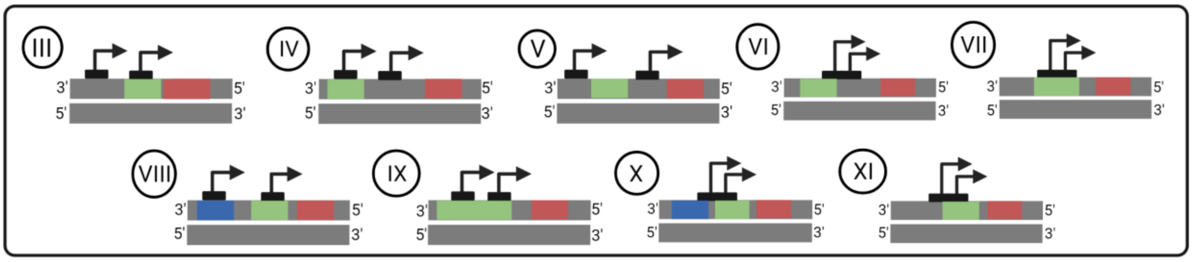


**Fig S1. Other arrangements of tandem promoters in *E. coli.*** Unlike the arrangements **I** and **II** in Fig. 1 in the main manuscript, the arrangements here (**III**-**XI**) allow for overlaps with or in between other gene(s). The red, green, and blue rectangles are DNA regions coding for RNA. These arrangements are not considered in this study. Figure created with BioRender.com.

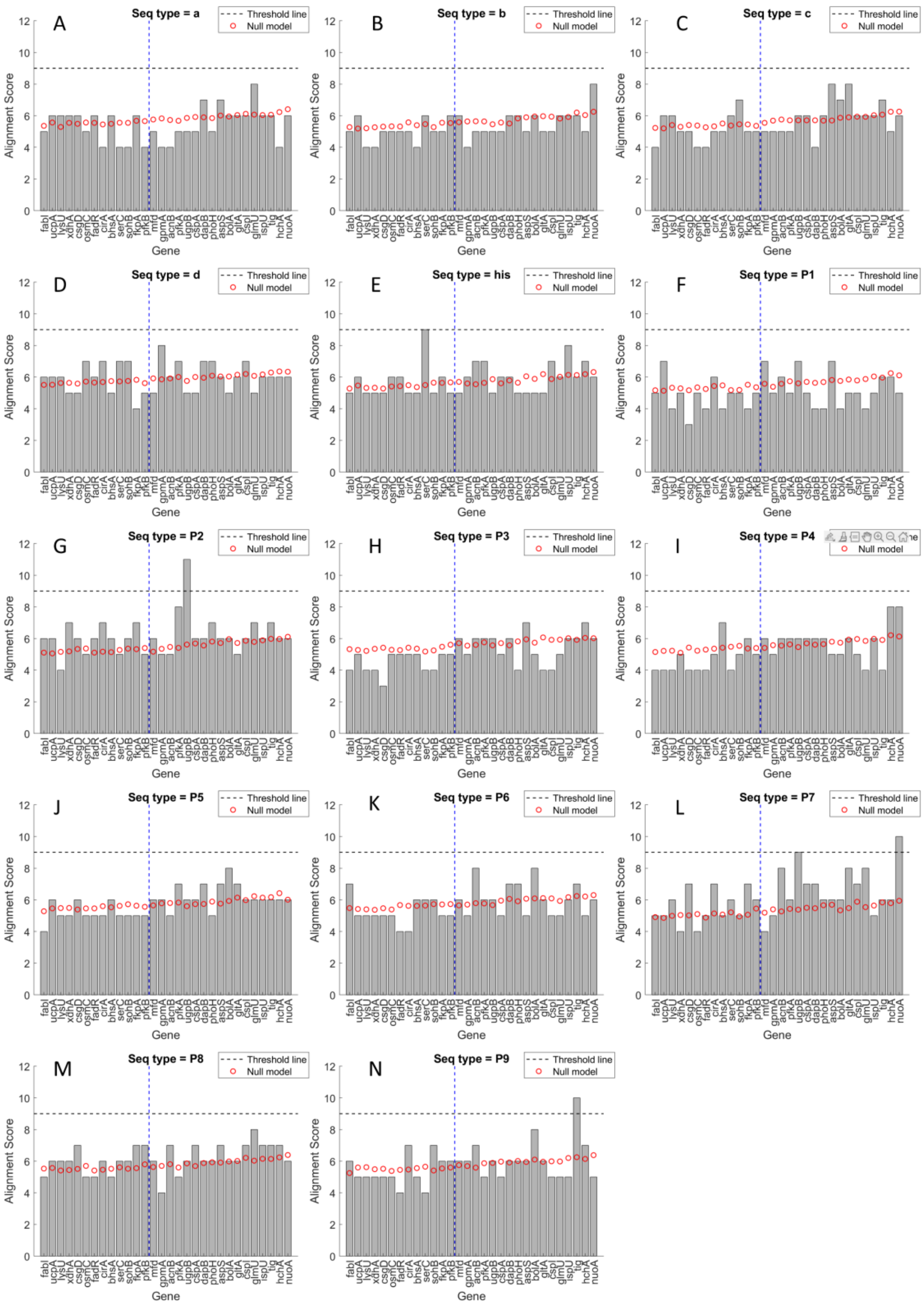


**Fig S2. Local alignment scores.** Local alignment scores between known pause sequences and the sequences in between the tandem promoter regions (grey bars). Also shown by red circles are the alignment scores between each pause sequence and randomly generated sequences with the same $d_{TSS}$ as the natural genes. The minimum alignment score to be considered significant is shown by a dashed black line. Finally, the blue vertical dashed line at $d_{TSS}$ = 35 bp shows the separation between genes subject to occlusion or not.

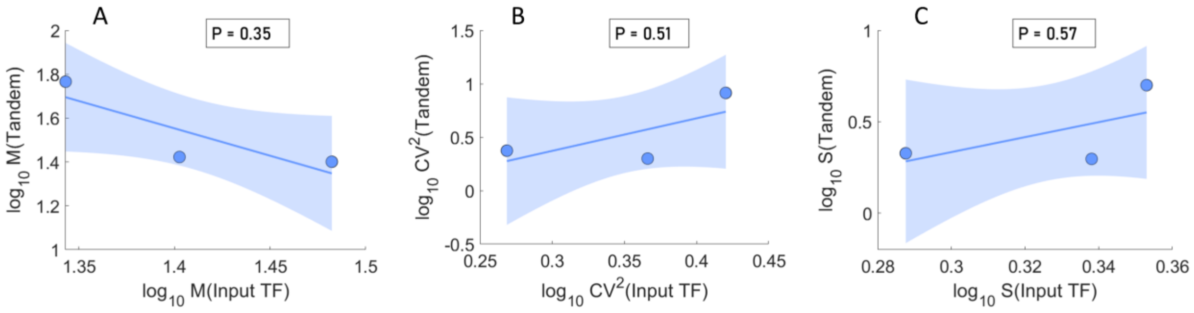


**Fig S3. Correlation of the moments of the single-cell protein numbers between genes and their input TFs.** Scatter plots between the moments of the single-cell protein numbers (in $log_{10}$ scale) of genes regulated by tandem promoters ('Tandem') and their input TFs. (A) Mean, (B) $CV^2$, and (C) Skewness. The blue line is the best linear fit, and its shadow is the standard error of the fit. The p-value, *P* is the probability that the slope of the line equals 0. If $P < 0.05$, there is a statistically significant correlation. The genes used in these results are listed in Table S5 in S3 Appendix. The axes differ widely in scales between the figures to facilitate visualization of the relationships.

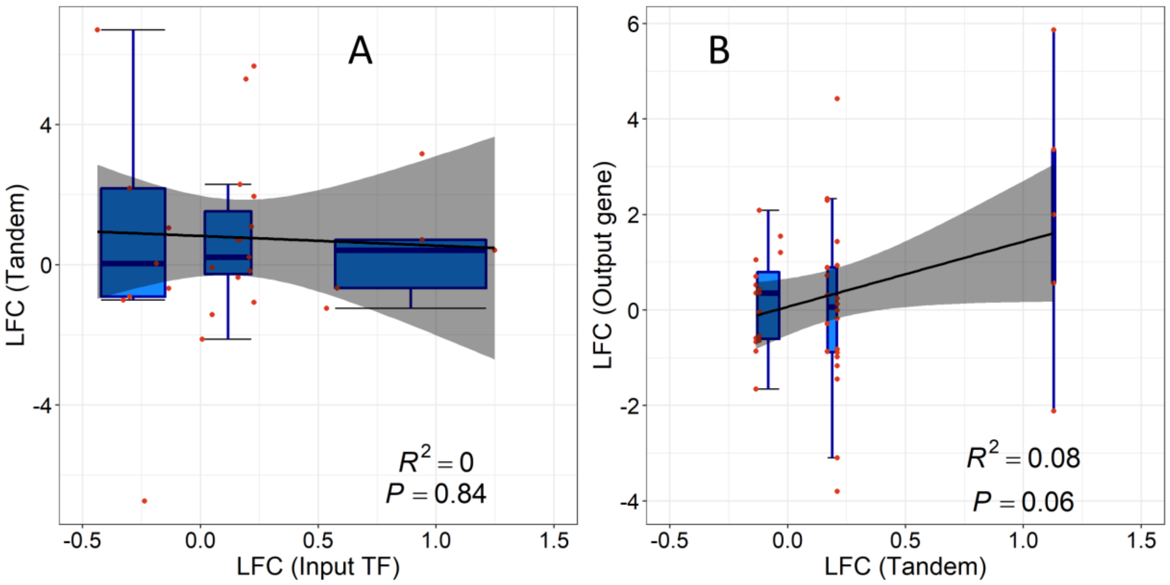


**Fig S4. Correlation of RNA fold changes of genes and their input TFs.** Correlation plots between the LFCs of the RNA numbers of genes controlled by tandem promoters with their input and output genes. (A) LFCs (from 0 to 20 min) of 29 genes expressing input TFs plotted against the corresponding LFCs (from 0 to 180 min) of the genes controlled by tandem promoters. (B) LFCs (from 0 to 20 min) of genes controlled by tandem promoters plotted against the corresponding LFCs of their output genes (from 0 to 180 min). A total of 43 TF-gene interactions were analysed. RNA-seq measurements described in section "RNA-seq Measurements and Analysis in S1 Appendix". The black line is the best linear fit and the grey shadow area is the standard error of the fit. The blue horizontal lines inside the boxes are the median, the top of the boxes are the 3rd quartile (Q3) and the bottom of the boxes are the first quartile (Q1). The error bars at the top and bottom range from (Q3+1.5*IQR) to (Q1-1.5*IQR), with an interquartile range: IQR = Q3 – Q1. The three box plots correspond to the data points with LFCs < 0, LFC between 0 and 0.5, and LFC > 0.5. Related to Table S5 and S6 in S3 Appendix.

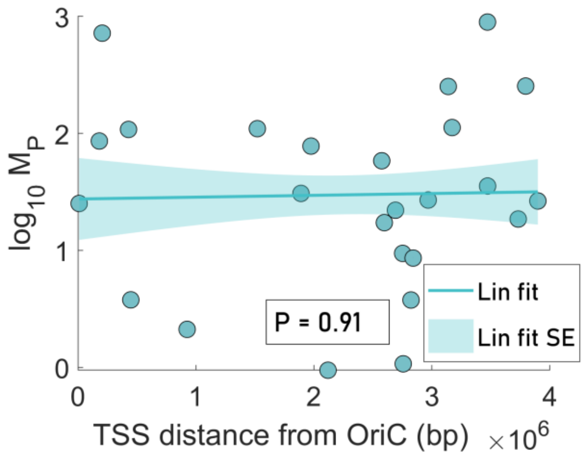


**Fig S5. Relationship between expression levels of the genes controlled by tandem promoters and the distance in nucleotides (bp) from the upstream promoter and the Oric region in the DNA.** Data from 25 genes for the 1X condition. Also shown in a linear fit and the corresponding 1 standard error of the fit (shadow area). The p-value, *P*, is the probability that the slope of the line equals 0.

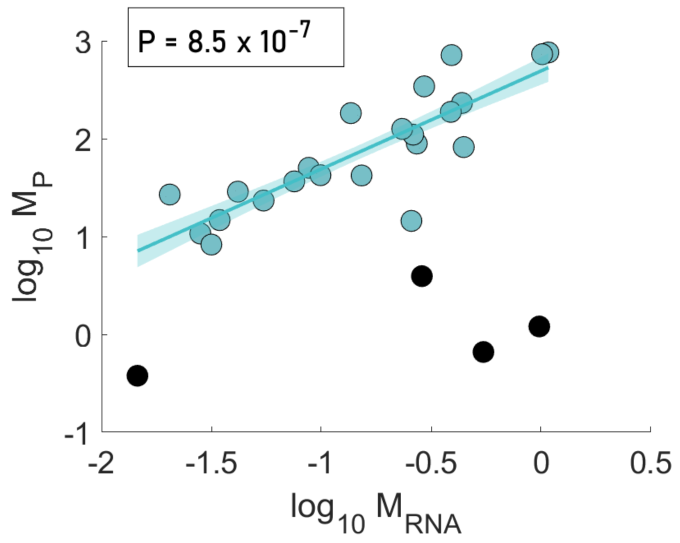


**Fig S6. Correlation plot between the mean single-cell RNA levels ($M_{RNA}$) and the mean single-cell protein numbers ($M_P$).** Both data are obtained from Ref. [28] in main manuscript and are processed to include only genes controlled by tandem promoters (classes **I** and **II**, Table S8 in S3 Appendix). The line is the best linear fit to the data, and its shadow area is the standard error of the fit. The p-value, *P* is the probability that the slope of the line equals 0. Since $P < 0.05$, we conclude that $M_{RNA}$ and $M_P$ are significantly correlated. The black balls correspond to 4 genes that were not considered when fitting the line, due to being outliners. In our own data, cells carrying these same 4 genes exhibited a fluorescence that was equal or lower than the cellular background fluorescence in either 1X or 0.5X media.

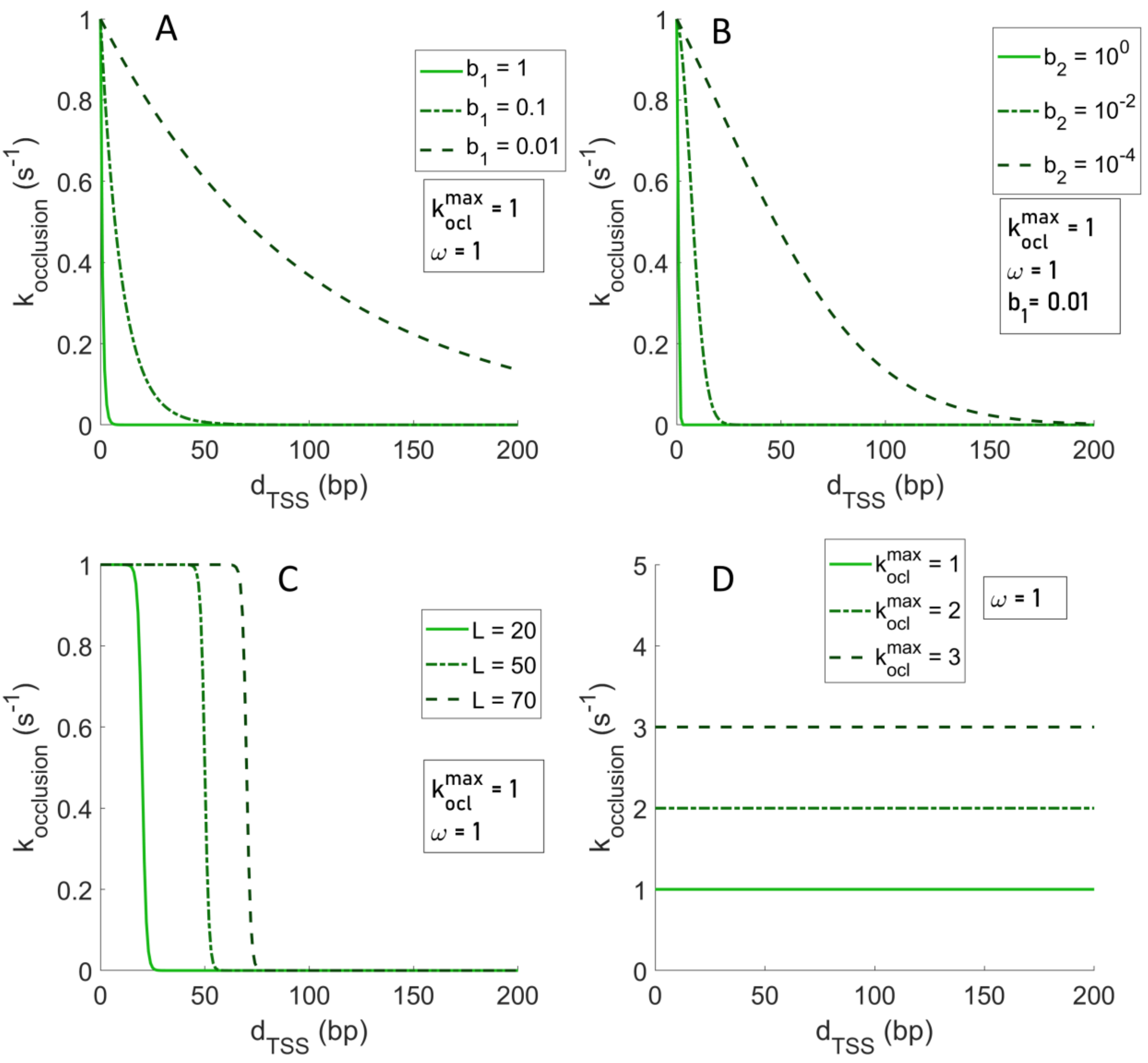


**Fig S7. Models of transcription interference.** Models of transcription interference between RNAPs in tandem promoters as a function of the $d_{TSS}$ between them. (A) 'Exponential 1' as a function for different values of '$b_1$'. (B) 'Exponential 2' as a function at different values of '$b_2$. (C) Continuous 'step-like' function for different values of '$L$' (which is the $d_{TSS}$ at which the step occurs). (D) Zero order polynomial for different values of $k_{ocl}^{max}$ . See Table 1 in the main manuscript for the definitions of these models and variables within.

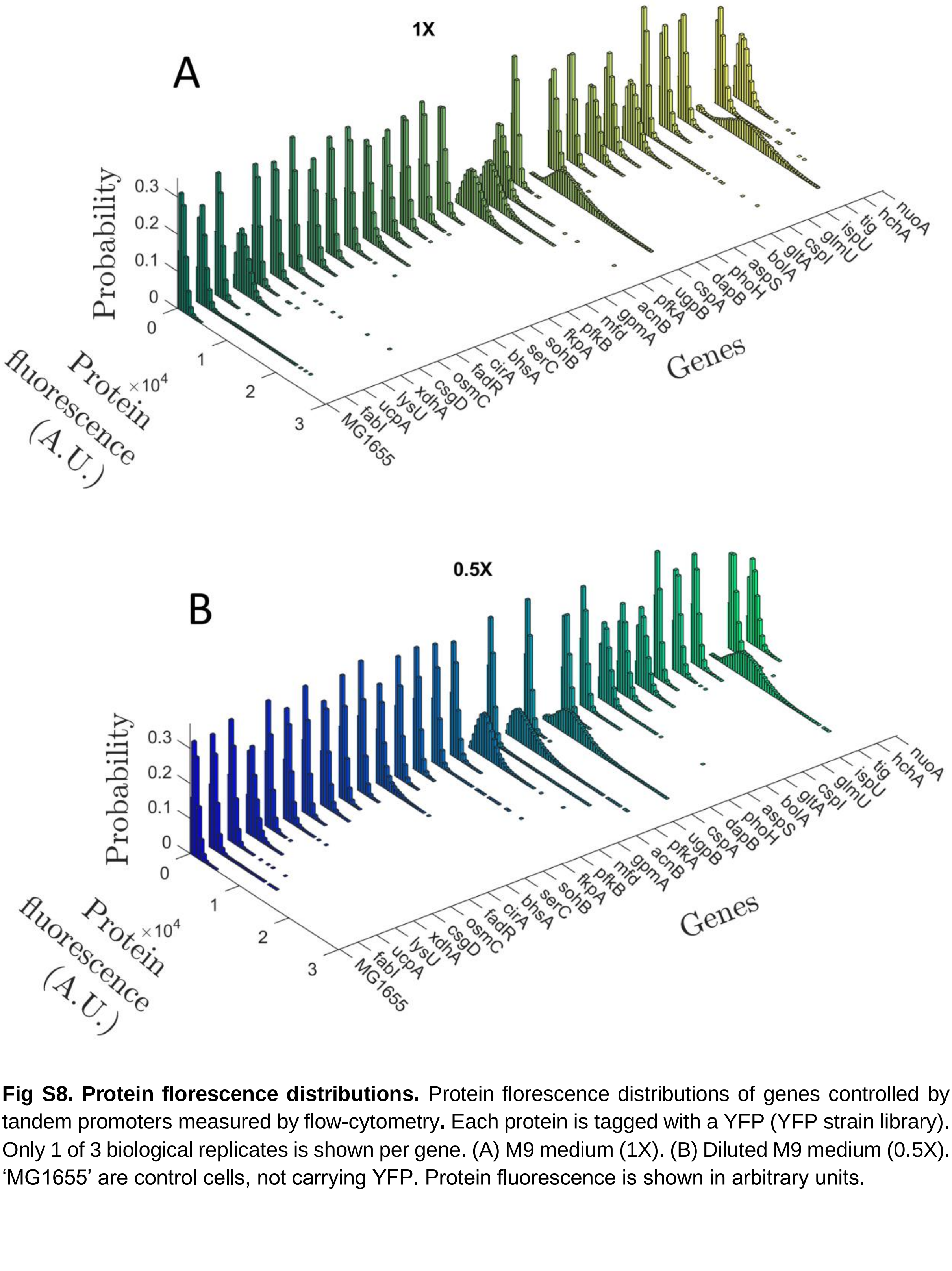


**Fig S8. Protein florescence distributions.** Protein florescence distributions of genes controlled by tandem promoters measured by flow-cytometry**.** Each protein is tagged with a YFP (YFP strain library). Only 1 of 3 biological replicates is shown per gene. (A) M9 medium (1X). (B) Diluted M9 medium (0.5X). ‘MG1655’ are control cells, not carrying YFP. Protein fluorescence is shown in arbitrary units.

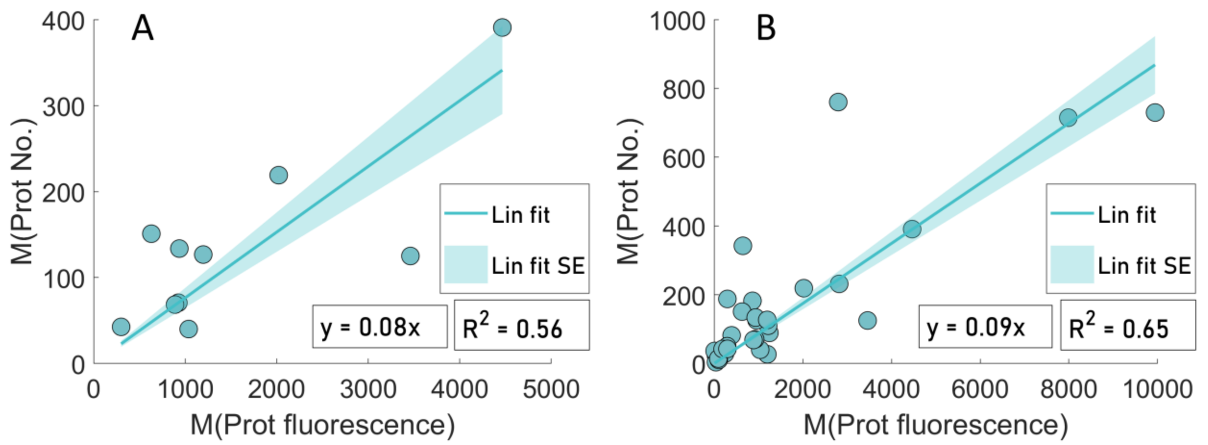


**Fig S9. Estimation of scaling factors using data from genes controlled by single promoters**. A) Mean single-cell protein fluorescence (own measurements of genes controlled by single promoters) plotted against the corresponding mean single-cell protein numbers reported in [28]. From the equation of the best fitting line without y-intercept (y-intercept = 0), we obtained a scaling factor, $s_f$, equal to 0.08. B) Same as (A) but the own measurements are of both single promoters and tandem promoters, merged. From the equation of the best fitting line without y-intercept (y-intercept = 0), we obtained a scaling factor, $s_f$, equal to 0.09.

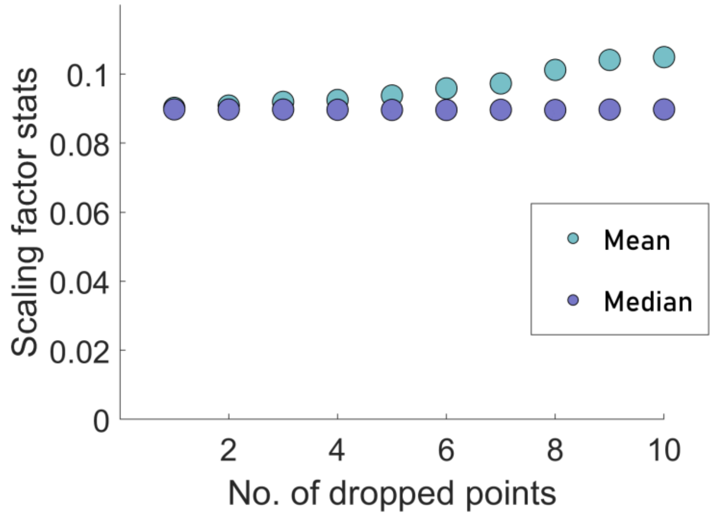


**Fig S10. Sensitivity test.** Mean and median of scaling factor varies as a function of number of data points randomly dropped.

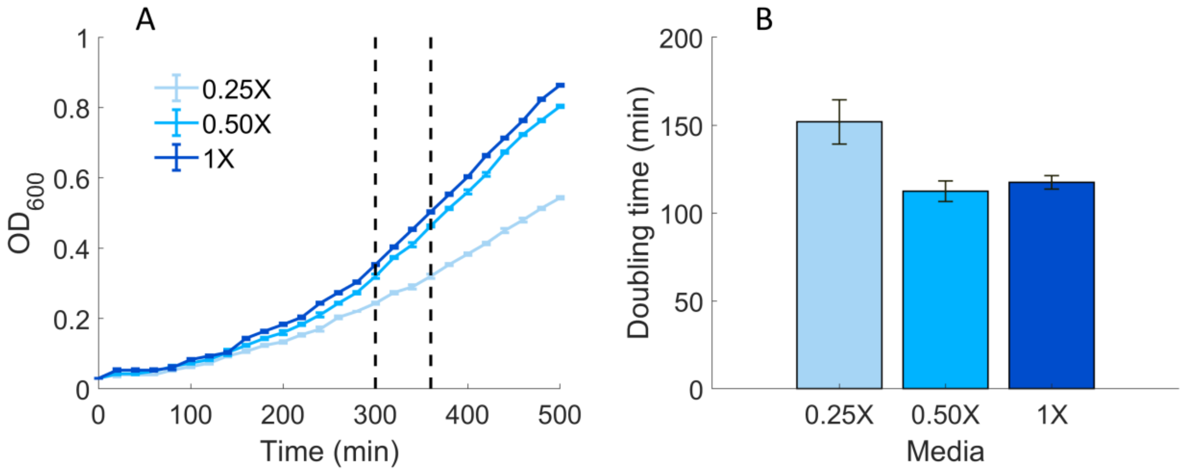


**Fig S11. Growth curves and doubling times.** A. Optical density ($OD_{600}$) curves of *E. coli* MG1655 cells grown in 0.25X,0.5X and 1X media (section 'Media and Chemicals' in S1 Appendix). B. From these curves, the doubling time was estimated to be ~112 min in 0.5X and ~118 min in 1X. We used 115 min doubling time in the models. The estimation is made using the formula

$$D = \frac{\ln(2)}{\ln\left(\frac{OD(t_2)}{OD(t_1)}\right)} \times (t_2 - t_1)$$

, with $t_2$ and $t_1$ being the end and start times (in minutes), respectively. They are marked by two vertical dashed black lines. The error bars denote the standard error of the mean. Ref. [28] in main manuscript reported ~150 min using 96 well-plates in the same conditions. The fact that we used culture tubes may explain the difference.

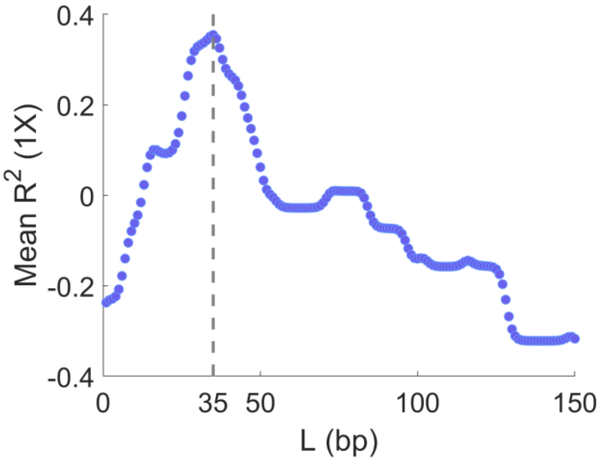


**Fig S12. Mean $R^2$ of the step interference model.** Mean $R^2$ of the step interference model to the 1X data in Figures 6A, 6B, and 6C, as a function of L ($d_{TSS}$ at which the step of the step function occurs). The Mean $R^2$ is visibly maximized at L = 35, which marked by a grey dashed line. Relates to Figure 6 in the main manuscript.

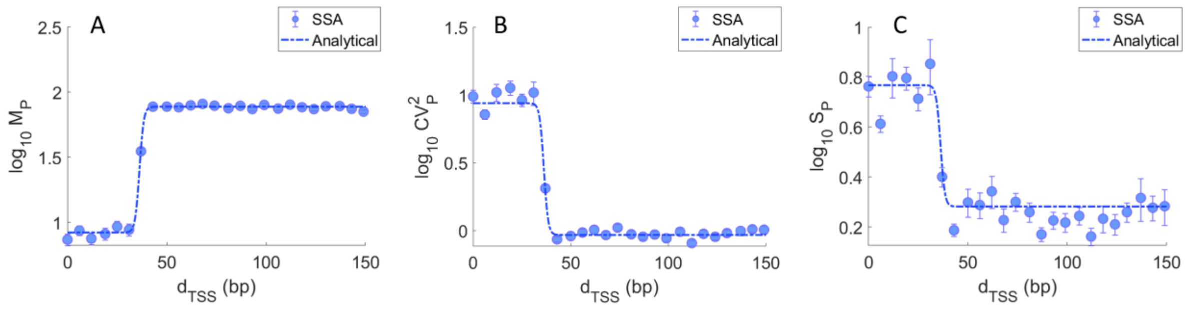


**Fig S13. Confronting the solutions of the analytical and stochastic model.** (A) $\mathrm{Log}_{10}$ of mean protein numbers, (B) $\log_{10}$ of $CV^2$ of protein numbers and (C) $\log_{10}$ of Skewness of protein numbers as a function of $d_{TSS}$. The blue line is the analytical solution of the step model. The blue dots are the mean results of stochastic simulations of the step model. The parameters used are shown in Tables 2 and 3 in the main manuscript. See Section 'Stochastic simulations for the step interference model' in S1 Appendix.

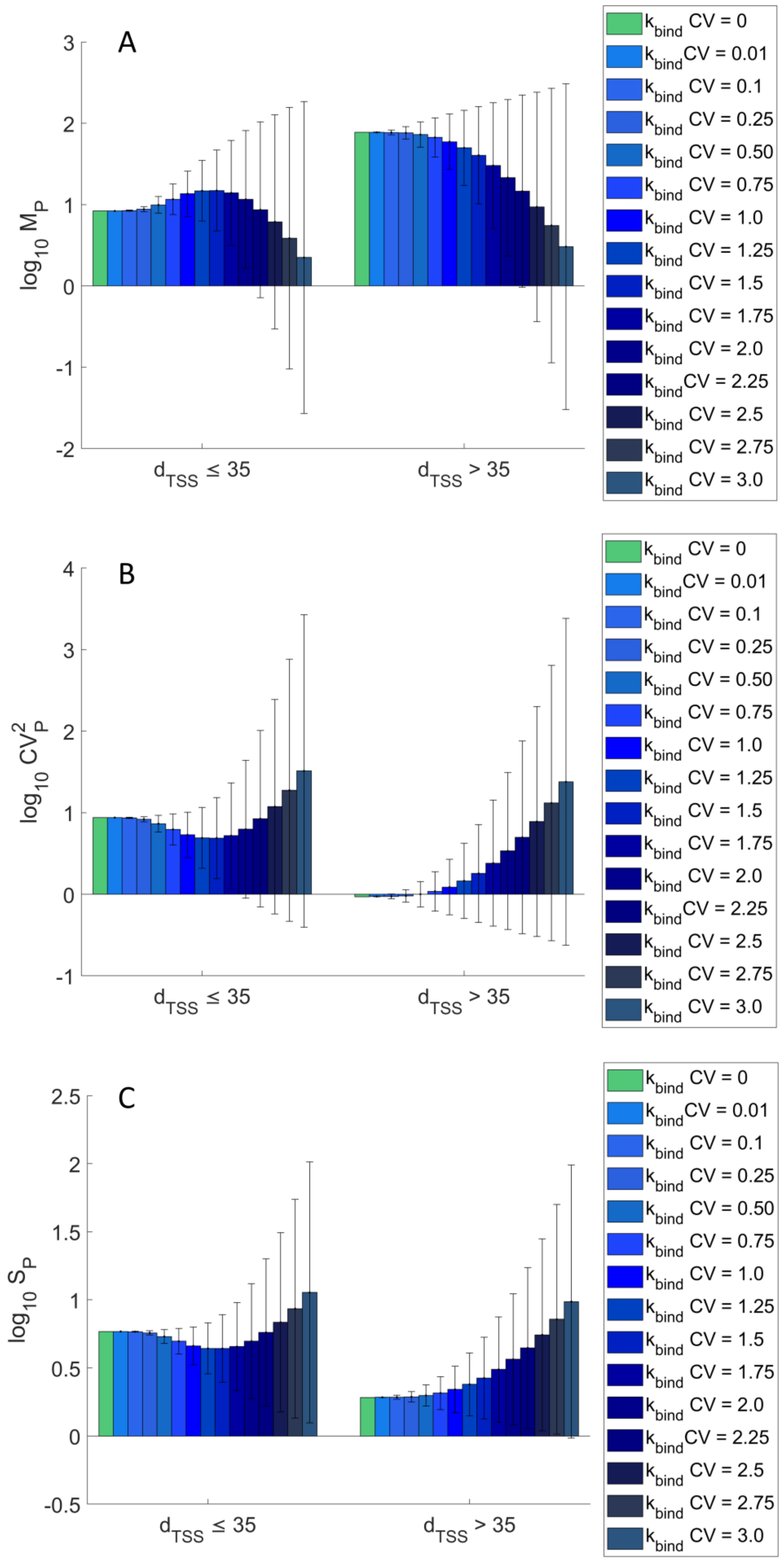
A
B
C
log10 MP
log10 CV2P
log10 SP
dTSS ≤ 35
dTSS > 35
kbind CV = 0
kbind CV = 0.01
kbind CV = 0.1
kbind CV = 0.25
kbind CV = 0.50
kbind CV = 0.75
kbind CV = 1.0
kbind CV = 1.25
kbind CV = 1.5
kbind CV = 1.75
kbind CV = 2.0
kbind CV = 2.25
kbind CV = 2.5
kbind CV = 2.75
kbind CV = 3.0

**Fig S14. Solutions of the analytical model for different levels of variability of $k_{bind} \cdot [R]$.** (Top) Mean, (Middle) CV[2] and (Bottom) S of single-cell protein numbers produced by tandem promoters when $d_{TSS}$ ≤ 35 (left) and $d_{TSS}$ > 35 (right). The green bar is the analytical solution with $CV\left(k_{bind} \cdot [R]\right) = 0$. The other bars are from analytical solutions for various degrees of variability of $k_{bind} \cdot [R]$ of each promoter.

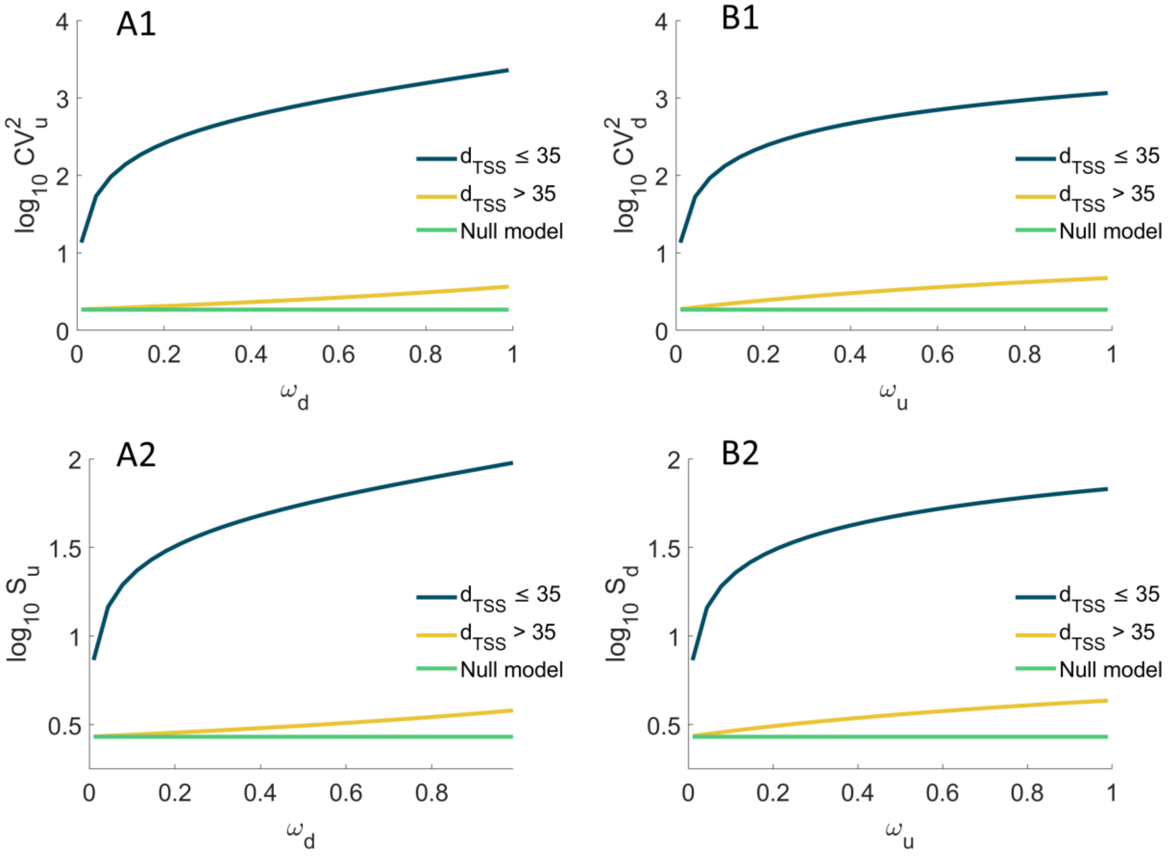


**Fig S15. Variability and skewness in single-cell protein numbers produced from an upstream and from a downstream promoter as a function of promoter occupancy of the other promoter.** $CV_P^2$ and $S_P$ of the single-cell distribution of the number of proteins produced (**A1 and A2**) by the upstream promoter alone, and (**B1 and B2**) by the downstream promoter alone. Results are shown as a function of the fraction of times that the upstream (0.01 ≤ $\omega_u$ ≤ 0.99) and the downstream (0.01 ≤ $\omega_d$ ≤ 0.99) promoter are occupied by RNAP. The null model is estimated by setting $k_{occlusion}$, $k_{sitting}$, and $\omega$ to zero.

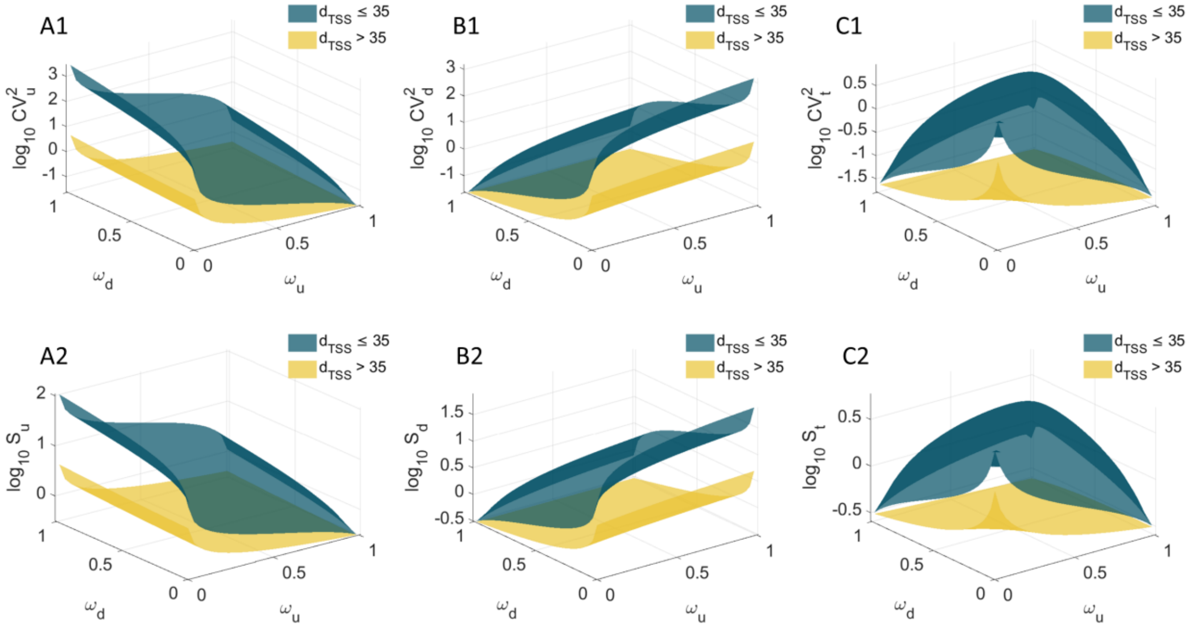


**Fig S16. Variability and skewness in single-cell protein numbers as a function of promoter occupancy.** Expected variability and skewness of the single cell distribution of protein numbers due to the activity of, respectively: (**A1** and **A2**) the upstream promoter alone, (**B1** and **B2**) the downstream promoter alone, and (**C1** and **C2**) both promoters. Shown is S of the distributions as a function of the fraction of times that the upstream ($0 \leq \omega_u \leq 1$) and the downstream ($0 \leq \omega_d \leq 1$) promoters are occupied by RNAP, when $d_{TSS} > 35$ (yellow) and $d_{TSS} \leq 35$ (dark green) bp.

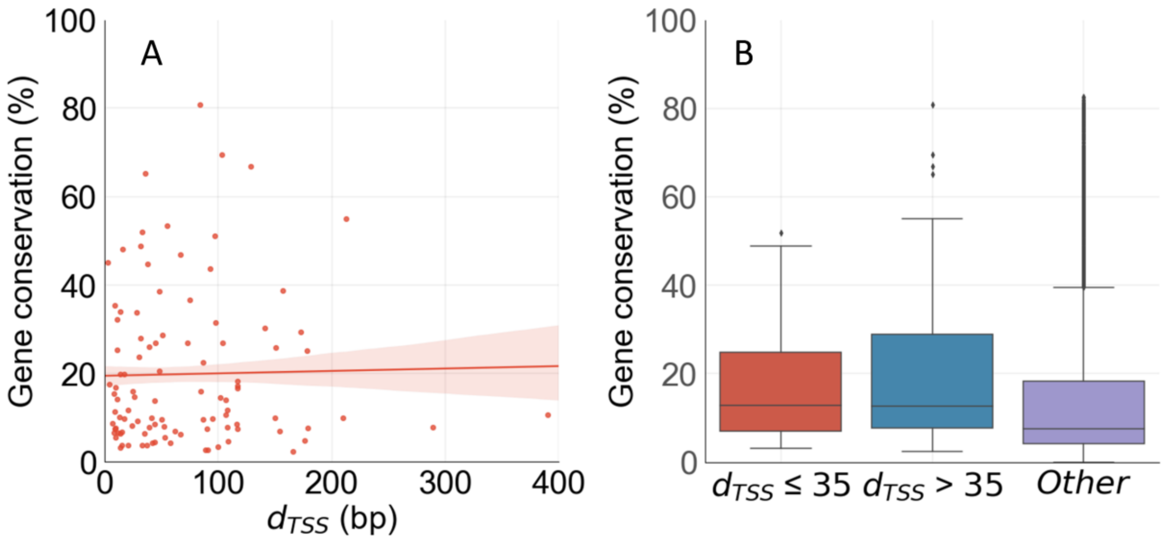


**Fig S17. Gene conservation levels**. (A) Correlation between $d_{TSS}$ (bp) of the pairs of tandem promoters and the evolutionary conservation level of the gene that they express. The line shown is the best linear fit to the data, and its shadow is the standard error of the fit. (B) Box plot of the gene conservation levels of the cohorts of genes with $d_{TSS} > 35$ and with $d_{TSS} \leq 35$, along with genes other than those in tandem formation. The horizontal black line inside each box marks the median, the top of the box shows the $3^{rd}$ quartile (Q3), and the bottom of the box shows the first quartile (Q1) of each gene cohort. The error bar above the box marks the range of values within (Q3+1.5*IQR), while the error bar below the bottom shows the range of values within (Q1-1.5*IQR). Here, IQR = Q3 – Q1.
