## Supporting Tables for "Analytical kinetic model of native tandem promoters in *E. coli*"

### S3 Appendix: Supporting Tables

**Table S1. List of genes controlled by tandem promoters.**

| S. No | Configuration (see Fig. 1 main manuscript) | Gene | Promoters (upstream/ downstream) | Distance between TSS's (bp) |
|---|---|---|---|---|
| **1** | I | aspS | aspSp1/aspSp | 84 |
| **2** | I | bolA | bolAp2/bolAp1 | 85 |
| **3** | I | cspI | cspIp/cspIp2 | 100 |
| **4** | I | glmU | glmUp2/glmUp1 | 103 |
| **5** | I | gltA | gltAp1/gltAp2 | 97 |
| **6** | I | hchA | hchAp2/hchAp | 150 |
| **7** | I | ispU | ispUp1/ispUp2 | 117 |
| **8** | I | tig | tigp1/tigp3 | 129 |
| **9** | I | nuoA | nuoAp1/nuoAp2 | 173 |
| **10** | II | acnB | acnBp/acnBp2 | 45 |
| **11** | II | bhsA | bhsAp9/bhsAp | 14 |
| **12** | II | cirA | cirAp2/cirAp1 | 13 |
| **13** | II | csgD | csgDp1/csgDp2 | 9 |
| **14** | II | cspA | cspAp1/cspAp2 | 51 |
| **15** | II | dapB | dapBp2/dapBp1 | 55 |
| **16** | II | fabI | fabIp/fabIp1 | 3 |
| **17** | II | fadR | fadRp/fadRp2 | 11 |
| **18** | II | fkpA | fkpAp1/fkpAp2 | 26 |
| **19** | II | gpmA | gpmAp2/gpmAp | 38 |
| **20** | II | lysU | lysUp1/lysUp2 | 8 |
| **21** | II | mfd | mfdp1/mfdp2 | 36 |
| **22** | II | osmC | osmCp1/osmCp2 | 10 |
| **23** | II | pfkA | pfkAp2/pfkAp1 | 48 |
| **24** | II | pfkB | pfkBp2/pfkBp1 | 28 |
| **25** | II | phoH | phoHp1/phoHp2 | 73 |
| **26** | II | serC | serCp2/serCp | 16 |
| **27** | II | sohB | sohBp1/sohBp2 | 17 |
| **28** | II | ucpA | ucpAp2/ucpAp1 | 7 |
| **29** | II | ugpB | ugpBp2/ugpBp1 | 48 |
| **30** | II | xdhA | xdhAp/xdhAp2 | 8 |

List of genes controlled by tandem promoters whose single-cell protein numbers were measured by flow-cytometry using cells of the YFP strain library. Also shown are their promoters in tandem formation, their configuration, and the distance in base pairs (bp) between their TSSs.

**Table S2. List of strains of the YFP strain library observed by flow-cytometry.**

| S. No. | Strain name | Genotype | Source |
|---|---|---|---|
| **1** | acnB [SX1900] | F-, acnB791-YFP(::cat), Δ(argF-lac)169, gal-490, Δ(modF-ybhJ)803, λ[cI857 Δ(cro-bioA)], IN(rrnD-rrnE)1, rph-1 | Yale CGSC (CGSC # 13455) |
| **2** | argP [SX1436] | F-, Δ(argF-lac)169, gal-490, Δ(modF-ybhJ)803, λ[cI857 Δ(cro-bioA)], argP794-YFP(::cat), IN(rrnD-rrnE)1, rph-1 | Yale CGSC (CGSC # 12991) |
| **3** | aspS [SX1044] | F-, Δ(argF-lac)169, gal-490, Δ(modF-ybhJ)803, λ[cI857 Δ(cro-bioA)], aspS793-YFP(::cat), IN(rrnD-rrnE)1, rph-1 | Yale CGSC (CGSC # 12599) |
| **4** | bhsA [SX1979] | F-, Δ(argF-lac)169, gal-490, Δ(modF-ybhJ)803, λ[cI857 Δ(cro-bioA)], bhsA791-YFP(::cat), IN(rrnD-rrnE)1, rph-1 | Yale CGSC (CGSC # 13534) |
| **5** | bolA [SX1087] | F-, Δ(argF-lac)169, bolA791-YFP(::cat), gal-490, Δ(modF-ybhJ)803, λ[cI857 Δ(cro-bioA)], IN(rrnD-rrnE)1, rph-1 | Yale CGSC (CGSC # 12642) |
| **6** | cirA [SX1509] | F-, Δ(argF-lac)169, gal-490, Δ(modF-ybhJ)803, λ[cI857 Δ(cro-bioA)], cirA791-YFP(::cat), IN(rrnD-rrnE)1, rph-1 | Yale CGSC (CGSC # 13064) |
| **7** | csgD [SX1465] | F-, Δ(argF-lac)169, gal-490, Δ(modF-ybhJ)803, λ[cI857 Δ(cro-bioA)], csgD791-YFP(::cat), IN(rrnD-rrnE)1, rph-1 | Yale CGSC (CGSC # 13020) |
| **8** | cspA [SX1097] | F-, Δ(argF-lac)169, gal-490, Δ(modF-ybhJ)803, λ[cI857 Δ(cro-bioA)], IN(rrnD-rrnE)1, cspA791-YFP(::cat), rph-1 | Yale CGSC (CGSC # 12652) |
| **9** | cspI [SX1106] | F-, Δ(argF-lac)169, gal-490, Δ(modF-ybhJ)803, λ[cI857 Δ(cro-bioA)], cspI797-YFP(::cat), IN(rrnD-rrnE)1, rph-1 | Yale CGSC (CGSC # 12661) |
| **10** | dapB [SX1910] | F-, dapB792-YFP(::cat), Δ(argF-lac)169, gal-490, Δ(modF-ybhJ)803, λ[cI857 Δ(cro-bioA)], IN(rrnD-rrnE)1, rph-1 | Yale CGSC (CGSC # 13465) |
| **11** | fabD [SX2002] | F-, Δ(argF-lac)169, gal-490, Δ(modF-ybhJ)803, λ[cI857 Δ(cro-bioA)], fabD793-YFP(::cat), IN(rrnD-rrnE)1, rph-1 | Yale CGSC (CGSC # 13557) |
| **12** | fabH [SX1474] | F-, Δ(argF-lac)169, gal-490, Δ(modF-ybhJ)803, λ[cI857 Δ(cro-bioA)], fabH795-YFP(::cat), IN(rrnD-rrnE)1, rph-1 | Yale CGSC (CGSC # 13029) |
| **13** | fabI [SX1038] | F-, Δ(argF-lac)169, gal-490, Δ(modF-ybhJ)803, λ[cI857 Δ(cro-bioA)], fabI796-YFP(::cat), IN(rrnD-rrnE)1, rph-1 | Yale CGSC (CGSC # 12593) |
| **14** | fadR [SX1521] | F-, Δ(argF-lac)169, gal-490, Δ(modF-ybhJ)803, λ[cI857 Δ(cro-bioA)], fadR795-YFP(::cat), IN(rrnD-rrnE)1, rph-1 | Yale CGSC (CGSC # 13076) |
| **15** | fkpA [SX2015] | F-, Δ(argF-lac)169, gal-490, Δ(modF-ybhJ)803, λ[cI857 Δ(cro-bioA)], IN(rrnD-rrnE)1, fkpA791-YFP(::cat), rph-1 | Yale CGSC (CGSC # 13570) |
| **16** | fur [SX1916] | F-, Δ(argF-lac)169, fur-791-YFP(::cat), gal-490, Δ(modF-ybhJ)803, λ[cI857 Δ(cro-bioA)], IN(rrnD-rrnE)1, rph-1 | Yale CGSC (CGSC # 13471) |

| | | | |
|---|---|---|---|
| **17** | glmU [SX1004] | F-, Δ(argF-lac)169, gal-490, Δ(modF-ybhJ)803, λ[cI857 Δ(cro-bioA)], IN(rrnD-rrnE)1, rph-1, glmU792-YFP(::cat) | Yale CGSC (CGSC # 12559) |
| **18** | gltA [SX1925] | F-, Δ(argF-lac)169, gltA791-YFP(::cat), gal-490, Δ(modF-ybhJ)803, λ[cI857 Δ(cro-bioA)], IN(rrnD-rrnE)1, rph-1 | Yale CGSC (CGSC # 13480) |
| **19** | gpmA [SX1553] | F-, Δ(argF-lac)169, gpmA791-YFP(::cat), gal-490, Δ(modF-ybhJ)803, λ[cI857 Δ(cro-bioA)], IN(rrnD-rrnE)1, rph-1 | Yale CGSC (CGSC # 13108) |
| **20** | hchA [SX1988] | F-, Δ(argF-lac)169, gal-490, Δ(modF-ybhJ)803, λ[cI857 Δ(cro-bioA)], hchA791-YFP(::cat), IN(rrnD-rrnE)1, rph-1 | Yale CGSC (CGSC # 13243) |
| **21** | ispU [SX1052] | F-, ispU796-YFP(::cat), Δ(argF-lac)169, gal-490, Δ(modF-ybhJ)803, λ[cI857 Δ(cro-bioA)], IN(rrnD-rrnE)1, rph-1 | Yale CGSC (CGSC # 12607) |
| **22** | lysU [SX1127] | F-, Δ(argF-lac)169, gal-490, Δ(modF-ybhJ)803, λ[cI857 Δ(cro-bioA)], IN(rrnD-rrnE)1, rph-1, lysU793-YFP(::cat) | Yale CGSC (CGSC # 12682) |
| **23** | mfd [SX1072] | F-, Δ(argF-lac)169, gal-490, Δ(modF-ybhJ)803, λ[cI857 Δ(cro-bioA)], mfd-791-YFP(::cat), IN(rrnD-rrnE)1, rph-1 | Yale CGSC (CGSC # 12627) |
| **24** | mreB [SX1466] | F-, Δ(argF-lac)169, gal-490, Δ(modF-ybhJ)803, λ[cI857 Δ(cro-bioA)], mreB791-YFP(::cat), IN(rrnD-rrnE)1, rph-1 | Yale CGSC (CGSC # 13021) |
| **25** | nagC [SX1561] | F-, Δ(argF-lac)169, nagC791-YFP(::cat), gal-490, Δ(modF-ybhJ)803, λ[cI857 Δ(cro-bioA)], IN(rrnD-rrnE)1, rph-1 | Yale CGSC (CGSC # 13116) |
| **26** | nlpA [SX1615] | F-, Δ(argF-lac)169, gal-490, Δ(modF-ybhJ)803, λ[cI857 Δ(cro-bioA)], IN(rrnD-rrnE)1, rph-1, nlpA791-YFP(::cat) | Yale CGSC (CGSC # 13170) |
| **27** | nuoA [SX1772] | F-, Δ(argF-lac)169, gal-490, Δ(modF-ybhJ)803, λ[cI857 Δ(cro-bioA)], nuoA791-YFP(::cat), IN(rrnD-rrnE)1, rph-1 | Yale CGSC (CGSC # 13327) |
| **28** | osmC [SX1758] | F-, Δ(argF-lac)169, gal-490, Δ(modF-ybhJ)803, λ[cI857 Δ(cro-bioA)], osmC791-YFP(::cat), IN(rrnD-rrnE)1, rph-1 | Yale CGSC (CGSC # 13313) |
| **29** | pepD [SX1530] | F-, pepD792-YFP(::cat), Δ(argF-lac)169, gal-490, Δ(modF-ybhJ)803, λ[cI857 Δ(cro-bioA)], IN(rrnD-rrnE)1, rph-1 | Yale CGSC (CGSC #13085) |
| **30** | pfkA [SX1349] | F-, Δ(argF-lac)169, gal-490, Δ(modF-ybhJ)803, λ[cI857 Δ(cro-bioA)], IN(rrnD-rrnE)1, rph-1, pfkA791-YFP(::cat) | Yale CGSC (CGSC # 12904) |
| **31** | pfkB [SX1761] | F-, Δ(argF-lac)169, gal-490, Δ(modF-ybhJ)803, λ[cI857 Δ(cro-bioA)], pfkB792-YFP(::cat), IN(rrnD-rrnE)1, rph-1 | Yale CGSC (CGSC # 13316) |
| **32** | phoH [SX1752] | F-, Δ(argF-lac)169, gal-490, Δ(modF-ybhJ)803, λ[cI857 Δ(cro-bioA)], phoH791-YFP(::cat), IN(rrnD-rrnE)1, rph-1 | Yale CGSC (CGSC # 13307) |
| **33** | serC [SX1390] | F-, Δ(argF-lac)169, gal-490, Δ(modF-ybhJ)803, λ[cI857 Δ(cro-bioA)], serC791-YFP(::cat), IN(rrnD-rrnE)1, rph-1 | Yale CGSC (CGSC # 12945) |
| **34** | sohB [SX1707] | F-, Δ(argF-lac)169, gal-490, Δ(modF-ybhJ)803, λ[cI857 Δ(cro-bioA)], sohB791-YFP(::cat), IN(rrnD-rrnE)1, rph-1 | Yale CGSC (CGSC # 13262) |
| **35** | tig [SX1140] | F-, Δ(argF-lac)169, tig-791-YFP(::cat), gal-490, Δ(modF-ybhJ)803, λ[cI857 Δ(cro-bioA)], IN(rrnD-rrnE)1, rph-1 | Yale CGSC (CGSC # 12695) |

| | | | |
|---|---|---|---|
| **36** | ucpA [SX1211] | F-, Δ(argF-lac)169, gal-490, Δ(modF-ybhJ)803, λ[cI857 Δ(cro-bioA)], ucpA791-YFP(::cat), IN(rrnD-rrnE)1, rph-1 | Yale CGSC (CGSC # 12766) |
| **37** | ugpB [SX1574] | F-, Δ(argF-lac)169, gal-490, Δ(modF-ybhJ)803, λ[cI857 Δ(cro-bioA)], IN(rrnD-rrnE)1, ugpB791-YFP(::cat), rph-1 | Yale CGSC (CGSC # 13129) |
| **38** | wrbA [SX1718] | F-, Δ(argF-lac)169, gal-490, Δ(modF-ybhJ)803, λ[cI857 Δ(cro-bioA)], wrbA791-YFP(::cat), IN(rrnD-rrnE)1, rph-1 | Yale CGSC (CGSC # 13273) |
| **39** | xdhA [SX1671] | F-, Δ(argF-lac)169, gal-490, Δ(modF-ybhJ)803, λ[cI857 Δ(cro-bioA)], xdhA792-YFP(::cat), IN(rrnD-rrnE)1, rph-1 | Yale CGSC (CGSC # 13226) |
| **40** | yccJ [SX1975] | F-, Δ(argF-lac)169, gal-490, Δ(modF-ybhJ)803, λ[cI857 Δ(cro-bioA)], yccJ791-YFP(::cat), IN(rrnD-rrnE)1, rph-1 | Yale CGSC (CGSC # 13530) |
| **41** | yccT [SX1368] | F-, Δ(argF-lac)169, gal-490, Δ(modF-ybhJ)803, λ[cI857 Δ(cro-bioA)], yccT792-YFP(::cat), IN(rrnD-rrnE)1, rph-1 | Yale CGSC (CGSC # 12923) |
| **42** | aldA [SX1901] | F-, Δ(argF-lac)169, gal-490, Δ(modF-ybhJ)803, λ[cI857 Δ(cro-bioA)], aldA791-YFP(::cat), IN(rrnD-rrnE)1, rph-1 | Yale CGSC (CGSC # 13456) |
| **43** | elaB [SX1695] | F-, Δ(argF-lac)169, gal-490, Δ(modF-ybhJ)803, λ[cI857 Δ(cro-bioA)], elaB792-YFP(::cat), IN(rrnD-rrnE)1, rph-1 | Yale CGSC (CGSC # 13250) |
| **44** | feoA [SX1781] | F-, Δ(argF-lac)169, gal-490, Δ(modF-ybhJ)803, λ[cI857 Δ(cro-bioA)], IN(rrnD-rrnE)1, feoA791-YFP(::cat), rph-1 | Yale CGSC (CGSC # 13336) |
| **45** | gcvT [SX1674] | F-, Δ(argF-lac)169, gal-490, Δ(modF-ybhJ)803, λ[cI857 Δ(cro-bioA)], gcvT792-YFP(::cat), IN(rrnD-rrnE)1, rph-1 | Yale CGSC (CGSC # 13229) |
| **46** | glpD [SX1550] | F-, Δ(argF-lac)169, gal-490, Δ(modF-ybhJ)803, λ[cI857 Δ(cro-bioA)], IN(rrnD-rrnE)1, glpD792-YFP(::cat), rph-1 | Yale CGSC (CGSC # 13105) |
| **47** | pepN [SX1519] | F-, Δ(argF-lac)169, gal-490, Δ(modF-ybhJ)803, λ[cI857 Δ(cro-bioA)], pepN794-YFP(::cat), IN(rrnD-rrnE)1, rph-1 | Yale CGSC (CGSC # 13074) |
| **48** | wrbA [SX1718] | F-, Δ(argF-lac)169, gal-490, Δ(modF-ybhJ)803, λ[cI857 Δ(cro-bioA)], wrbA791-YFP(::cat), IN(rrnD-rrnE)1, rph-1 | Yale CGSC (CGSC # 13273) |
| **49** | ybeL [SX1822] | F-, Δ(argF-lac)169, ybeL794-YFP(::cat), gal-490, Δ(modF-ybhJ)803, λ[cI857 Δ(cro-bioA)], IN(rrnD-rrnE)1, rph-1 | Yale CGSC (CGSC # 13377) |
| **50** | ydfG [SX1986] | F-, Δ(argF-lac)169, gal-490, Δ(modF-ybhJ)803, λ[cI857 Δ(cro-bioA)], ydfG791-YFP(::cat), IN(rrnD-rrnE)1, rph-1 | Yale CGSC (CGSC # 13541) |
| **51** | yjbQ [SX1859] | F-, Δ(argF-lac)169, gal-490, Δ(modF-ybhJ)803, λ[cI857 Δ(cro-bioA)], IN(rrnD-rrnE)1, rph-1, yjbQ792-YFP(::cat) | Yale CGSC (CGSC # 13414) |

**Table S3. Average 'network' properties of genes with 1 or more TFs.**

| **Network properties** | **Genes controlled by tandem promoters with $d_{TSS}$ ≤ 35** | **Genes controlled by tandem promoters with $d_{TSS}$ > 35** | **All promoters of genes with 1 or more TF interactions** |
|---|---|---|---|

| | Mean ± SEM | Random set from all genes Mean ±SEM (p-value) | Mean ± SEM | Random set from all genes Mean ±SEM (p-value) | Mean ± SEM |
|---|---|---|---|---|---|
| **Average Shortest PathLength** | 0.31 ± 0.16 | 0.17 ± 0.11 (0.23) | 0.13 ± 0.05 | 0.17±0.08 (0.60) | 0.17 ±0.01 |
| **Clustering Coefficient** | 0.09 ± 0.03 | 0.11 ± 0.03 (0.68) | 0.10 ± 0.03 | 0.11 ± 0.03 (0.62) | 0.11 ± 4.34×$10^{-3}$ |
| **Eccentricity** | 0.56±0.31 | 0.25 ± 0.20 (0.22) | 0.15 ± 0.06 | 0.26 ± 0.16 (0.73) | 0.26 ± 0.03 |
| **Edge Count** | 5±1.64 | 4.64 ± 3.4 (0. 33) | 3.3 ± 0.83 | 4.64 ± 2.73 (0.67) | 4.63 ± 0.43 |
| **Indegree** | 2.33±0.48 | 2.32 ± 0.34 (0.52) | 2.02 ± 0.17 | 2.31 ± 0.27(0.83) | 2.32 ± 0.04 |
| **Neighborhood Connectivity** | 161.76 ± 29.09 | 131.95 ± 21.74 (0.20) | 134.63 ± 15.1 | 131.87 ± 17.36 (0. 44) | 131.91 ± 2.74 |
| **Outdegree** | 2.66 ± 1.34 | 2.33 ± 3.4 (0.30) | 1.28 ± 0.83 | 2.31 ± 2.71 (0. 59) | 2.32 ± 0.43 |

Shown are the network properties for genes controlled by tandem promoters at a distance $d_{TSS} \leq 35$ bp and at a distance $d_{TSS} > 35$ bp. For comparison, we show the same properties, when averaged from all genes of *E. coli*'s TF network. Genes without TF's are not considered. Note that all p-values are larger than 0.05.

**Table S4: Genes controlled by tandem promoters without input TFs.**

| S. No. | Gene | Availability in the YFP strain library |
|---|---|---|
| **1** | ampH | |
| **2** | ansP | |
| **3** | aroK | |
| **4** | aspS | ✓ |
| **5** | bepA | |
| **6** | cfa | |
| **7** | cobU | |
| **8** | crfC | |
| **9** | degQ | |
| **10** | fkpA | ✓ |
| **11** | ispU | ✓ |
| **12** | lpp | |
| **13** | mepS | |
| **14** | mfd | ✓ |
| **15** | narU | |

| | | |
|---|---|---|
| **16** | opgG | |
| **17** | panD | |
| **18** | pfkB | ✓ |
| **19** | serW | |
| **20** | tig | ✓ |
| **21** | ucpA | ✓ |
| **22** | xapR | |
| **23** | ybgI | |
| **24** | ygiM | |
| **25** | yheO | |
| **26** | yobF | |

Genes controlled by tandem promoters without input TFs. Those genes whose proteins are tagged with YFP in the YFP strain library are marked with the symbol '✓'.

**Table S5. Genes controlled by tandem promoters regulated by one and only one input TF.**

| | **Tandem promoter's genes** | **Availability in YFP strain library** | **Input TF** | **Availability in YFP strain library** |
|---|---|---|---|---|
| **1** | argR | | argR | |
| **2** | cvpA | | purR | |
| **3** | cysK | | cysB | ✓ |
| **4** | dapB | ✓ | argP | ✓ |
| **5** | fabI | ✓ | fadR | ✓ |
| **6** | fadR | ✓ | fadR | ✓ |
| **7** | fliL | | flhdC | |
| **8** | ftnB | | cpxR | ✓ |
| **9** | glgS | | crp | |
| **10** | glk | | cra | |
| **11** | glmU | ✓ | nagC | ✓ |
| **12** | gpmA | ✓ | fur | ✓ |
| **13** | hchA | ✓ | h-ns | |
| **14** | ibaG | | mlrA | ✓ |
| **15** | iraP | | csgD | ✓ |
| **16** | leuL | | leuO | |
| **17** | livK | | lrp | |
| **18** | lysU | ✓ | lrp | |
| **19** | mqsR | | mqsA | |
| **20** | ompA | | crp | |
| **21** | ompX | | fnr | |
| **22** | osmB | | rcsB | ✓ |
| **23** | pfkA | ✓ | cra | |

| | | | | |
|---|---|---|---|---|
| **24** | phoH | ✓ | phoB | |
| **25** | potF | | ntrC | |
| **26** | slyB | | phoP | |
| **27** | sohB | ✓ | crp | |
| **28** | wza | | rcsaB | |
| **29** | xdhA | ✓ | fnr | |
| **30** | ydbK | ✓ | soxS | |
| **31** | yeaG | | ntrc | |
| **32** | yhbT | | csgD | ✓ |
| **33** | yqjA | | cpxR | ✓ |

When the proteins of these genes and of their input TFs can be measured using strains of the YFP strain library, they are flagged with the symbol '✓'.

**Table S6. Genes controlled by, and only by, a TF expressed by tandem promoters.**

| | **Genes controlled by tandem promoters** | **Availability in YFP strain library** | **Genes regulated by the protein expressed by the gene controlled by tandem promoters** | **Availability in YFP strain library** |
|---|---|---|---|---|
| **1** | argR | | argA | ✓ |
| **2** | argR | | argB | |
| **3** | argR | | argC | |
| **4** | argR | | argE | ✓ |
| **5** | argR | | argF | |
| **6** | argR | | argH | |
| **7** | argR | | argI | |
| **8** | argR | | argR | |
| **9** | argR | | artI | |
| **10** | argR | | artJ | |
| **11** | argR | | artM | |
| **12** | argR | | artP | ✓ |
| **13** | argR | | artQ | |
| **14** | argR | | lysO | |
| **15** | bolA | ✓ | ampC | |
| **16** | bolA | ✓ | dacC | |
| **17** | bolA | ✓ | mreB | ✓ |
| **18** | bolA | ✓ | mreC | |
| **19** | bolA | ✓ | mreD | |
| **20** | csgD | ✓ | dgcC | |
| **21** | csgD | ✓ | iraP | |
| **22** | csgD | ✓ | nlpA | ✓ |
| **23** | csgD | ✓ | pepD | ✓ |
| **24** | csgD | ✓ | wrbA | ✓ |
| **25** | csgD | ✓ | yccJ | ✓ |
| **26** | csgD | ✓ | yccT | ✓ |

| | | | | |
|---|---|---|---|---|
| **27** | csgD | √ | yhbS | |
| **28** | csgD | √ | yhbT | |
| **29** | evgA | | frc | |
| **30** | evgA | | oxc | √ |
| **31** | evgA | | yegR | √ |
| **32** | evgA | | yegZ | |
| **33** | evgA | | yfdE | |
| **34** | evgA | | yfdV | |
| **35** | evgA | | yfdX | |
| **36** | fadR | √ | accA | |
| **37** | fadR | √ | accD | |
| **38** | fadR | √ | fabD | √ |
| **39** | fadR | √ | fabG | |
| **40** | fadR | √ | fabH | √ |
| **41** | fadR | √ | fabI | √ |
| **42** | fadR | √ | fadM | |
| **43** | fadR | √ | fadR | √ |
| **44** | xapR | | xapA | |
| **45** | xapR | | xapB | |

**Table S7. Protein levels and $d_{TSS}$ of 10 genes as measured by Microscopy and Image Analysis**.

| **Gene** | **TSS distance ($d_{TSS}$)** | **Mean single-cell protein level (Microscopy)** |
|---|---|---|
| xdhA | 8 | 0.04 |
| csgD | 9 | 0.64 |
| serC | 16 | 0.24 |
| sohB | 17 | 0.37 |
| pfkA | 48 | 2.8 |
| dapB | 55 | 0.57 |
| aspS | 84 | 1.72 |
| gltA | 97 | 3.02 |
| hchA | 150 | 0.74 |
| nuoA | 173 | 2.04 |

Related to Fig 4C in the main manuscript.

**Table S8. Number of genes controlled by a pair of tandem promoters in each configuration.**

| **Configuration** | **Number (in RegulonDB)** | **Present in the YFP strain library (measured here by flow-cytometry)** |
|---|---|---|
| **I** | 40 | 9(9) |
| **II** | 62 | 21(21) |
| **III** | 7 | 3 |

| | | |
|---|---|---|
| **IV** | 4 | 1 |
| **V** | 6 | 2 |
| **VI** | 0 | 0 |
| **VII** | 3 | 1 |
| **VIII** | 2 | 2 |
| **IX** | 4 | 1 |
| **X** | 0 | 0 |
| **XI** | 9 | 2 |
| **Other** | 6 | 0 |

Related to Fig 1in the main manuscript and Fig S1 in S2 Appendix.

**Table S9. Coefficient of variation, CV, of the gamma distribution.**

| **CV** **(**$k_{bind}\cdot[R]$**)** | $Mean\left(abs\left(k_{bind}^{u}\cdot[R]-k_{bind}^{d}\cdot[R]\right)\right)$ | $Mean\left(\frac{abs\left(k_{bind}^{u}\cdot[R]-k_{bind}^{d}\cdot[R]\right)}{k_{bind}^{u}\cdot[R]}\right)\times 100\%$ |
|---|---|---|
| 0.01 | $7.52 \times 10^{1}$ | 1.14 % |
| 0.1 | $7.64 \times 10^{-4}$ | $1.16 \times 10^{1}$ % |
| 0.25 | $1.86 \times 10^{-3}$ | $2.98 \times 10^{1}$ % |
| 0.5 | $3.63 \times 10^{-3}$ | $7.33 \times 10^{1}$ % |
| 0.75 | $5.27 \times 10^{-3}$ | $1.99 \times 10^{2}$ % |
| 1 | $6.62 \times 10^{-3}$ | $2.05 \times 10^{3}$ % |
| 1.25 | $7.81 \times 10^{-3}$ | $5.15 \times 10^{4}$ % |
| 1.5 | $8.66 \times 10^{-3}$ | $1.95 \times 10^{7}$ % |
| 1.75 | $9.41 \times 10^{-3}$ | $6.19 \times 10^{12}$ % |
| 2.0 | $9.89 \times 10^{-3}$ | $1.48 \times 10^{15}$ % |
| 2.25 | $1.04 \times 10^{-2}$ | $1.77 \times 10^{17}$ % |
| 2.5 | $1.10 \times 10^{-2}$ | $6.60 \times 10^{18}$ % |
| 2.75 | $1.12 \times 10^{-2}$ | $4.00 \times 10^{24}$ % |
| 3.0 | $1.20 \times 10^{-2}$ | $6.03 \times 10^{30}$ % |

Coefficient of variation, CV, of the gamma distribution from which $k_{bind}\cdot[R]$ of each promoter in tandem configuration is sampled from. Also shown is the resulting expected mean absolute difference in $k_{bind}\cdot[R]$ between the upstream and downstream promoters. Furthermore, the last column shows how much larger (in percentage) is one of the $k_{bind}\cdot[R]$ values compared to the other.

**Table S10. Location of the tandem promoters** relative to the oriC.

| Genes controlled by tandem promoters | Distance between the upstream TSS and the oriC |
|---|---|
| aspS | 1975043 |
| bolA | 3471395 |
| cspI | 2286932 |
| glmU | 10418 |
| gltA | 3170977 |
| hchA | 1890114 |
| ispU | 3730960 |
| nuoA | 1520409 |
| tig | 3470751 |
| acnB | 3794225 |
| bhsA | 2756725 |
| cirA | 1678802 |
| csgD | 2822400 |
| cspA | 205855 |
| dapB | 3897456 |
| fabI | 2574623 |
| fadR | 2690839 |
| fkpA | 448219 |
| gpmA | 3138074 |
| lysU | 428830 |
| mfd | 2751716 |
| osmC | 2369148 |
| pfkA | 181499 |
| pfkB | 2119421 |
| phoH | 2840879 |
| serC | 2968165 |
| sohB | 2596460 |
| ucpA | 1381073 |
| ugpB | 333318 |
| xdhA | 925487 |
