## Supporting Results for "Analytical kinetic model of native tandem promoters in *E. coli*"

### S4 Appendix: Supplementary Results

#### Pause sequences

We investigated if the nucleotide sequence of and in between the natural tandem promoters is coding for specific sequences known to perturb RNAP elongation. There are several events that compete with stepwise elongation. However, arrest, misincorporation and editing, pyrophosphorolysis, and premature termination are too rare in optimal growth conditions (rate constants listed in [1]) to be influential in several genes, and/or are not sequence dependent. Only sequences known to enhance transcriptional pausing [2] could fit both of these requirements. In *E. coli*, the mean rate of non-sequence specific pauses is 1 per 100 base pairs. These last 3 s on average [3-4]. However, a few sequences can enhance pausing frequency and/or duration (up to 15 or more seconds) [5] via various mechanically processes, which explains their variability in half-life and frequency of occurrence. For example, '*his*' pauses occur when the assembling RNA forms a hairpin-like loop, while '*ops*' pauses do not require it. Likely because of it, *his* pauses have longer half-life [6]. We searched in (and in between) the sequences of the 102 pairs of tandem promoters for the 14 sequences (each 12 nucleotides long) known to enhance pausing [7] (section 'Sequences prone to causes transcriptional pauses' in S1 Appendix) but found none. Thus, sequence-dependent transcriptional pausing should not be a common phenomenon in the tandem promoters of arrangements I and II. Even when allowing for 3 or less mistakes (sequence gaps, misalignments, duplicates, etc.), we only found 5 matches in the 30 of the 102 tandem promoter pairs studied with protein measurements below (Fig S2 in S2 Appendix, note the 5 bars crossing the threshold).

#### Over-representation test

We performed an over-representation test to search for biological functions (as defined in [8, 9] that are overrepresented by genes controlled by tandem promoters (using PANTHER 14 [10]). While based on a Fisher test, some biological processes appear to be overrepresented in our genes of interest (e.g., regulation of catabolic processes), none of them were significant to 'FDR correction' (FDR < 0.05, [10]. As such, we failed to identify a biological process significantly associated to genes controlled by tandem promoters (S5 Table).

#### Input-output transcription factor relationships

From time-lapse RNA-seq data, we assessed if the 102 genes controlled by tandem promoters (arrangements I and II, Fig 1) are affected by their input TFs. To facilitate this, we considered only those that have one and only input TF. I.e., we did not consider the 26 genes that do not have known input TFs (Table S4 in S3 Appendix), neither the 43 genes that have more than one input TF, making the detection of input-output relationships problematic. As such, of the 102, we considered only 33 genes (Table S5 in S3 Appendix). In these, we did not observe influences from input TFs (Figs S3 and S4A in

S2 Appendix). Finally, and similarly, we observed genes whose only input TF is expressed by tandem promoters (Table S6 in S3 Appendix). Again, we found no correlation (Fig S4B in S2 Appendix). Note that, while we did not find influences from TF interactions in the conditions of our measurements, we expect these interactions to become active in other conditions (e.g., stress conditions).

### Proteins with membrane-related positionings

From RegulonDB [11], of the 30 genes measured by flow-cytometry (Table S1 in S3 Appendix), only 3 are known to be related to membrane transportation and binding: bhsA, which is an outer membrane protein that is involved in copper permeability, stress resistance and biofilm formation, cirA, which is also an outer membrane transporter, and ugpB which is a periplasmic binding protein. Such membrane localizations could affect their quantification by YFP fusion, potentially by enhancing effects from avidity due to weakened diffusion.

However, none of these proteins significantly affect our results since, first, cirA and ugpB were removed from our analysis of the 1X condition, after preprocessing (gating, background subtraction and protein number conversion) (marked in red in S6 Table). Meanwhile, all three genes were removed from our analysis of the 0.5X condition after preprocessing (marked in red in S6 Table). Specifically, their removal was due to lack of expression above background autofluorescence.

### Relationship with the OriC region

From EcoCyc [12], the OriC region has a length of 232 base pairs and is located in positions 3 925 744 and 3 925 975 in the DNA of *E. coli*. We calculated the shortest distance between the TSS of the upstream promoter and the Oric region. These positions in the DNA are shown in Table S10 in the S3 Appendix. Meanwhile, the corresponding protein expression levels of these genes in the 1X condition are shown in the S6 Table. Finally, we show a Fig S5 in the S2 Appendix of these distances from OriC plotted again $\log_{10} M_p$ which shows that the two quantities do not correlate statistically.

### Regulation by H-NS

From RegulonDB [11], we investigated how many of the 102 genes controlled by tandem promoters (arrangements I and II) and how many of 30 of them observed by flow-cytometry are expected to be regulated by H-NS.

Of the 102 genes, 14 are regulated by H-NS (14%). Meanwhile, of the 30 genes, 5 are regulated by H-NS (17%). From this, we conclude that H-NS is not consistently a master regulator of these genes.

Nevertheless, of 4698 genes in E. coli, only 4 % are regulated by H-NS. This is significantly lower than in the case of the genes controlled by tandem promoters (p-value < 0.05 based on a Fisher test). As such, one could argue that H-NS regulation does occur higher than expected by chance. Future studies of the dynamics of those genes during environmental changes may thus be of interest.
